# A functional screen uncovers circular RNAs regulating excitatory synaptogenesis in hippocampal neurons

**DOI:** 10.1101/2024.04.09.588686

**Authors:** Darren Kelly, Silvia Bicker, Jochen Winterer, Prakruti Nanda, Pierre-Luc Germain, Christoph Dieterich, Gerhard Schratt

## Abstract

Circular RNAs (circRNAs) are an expanding class of largely unexplored RNAs which are prominently enriched in the mammalian brain. Here, we systematically interrogated their role in excitatory synaptogenesis of rat hippocampal neurons using RNA interference. Thereby, we identified seven circRNAs as negative regulators of excitatory synapse formation, many of which contain high-affinity microRNA binding sites. Knockdown of one of these candidates, circRERE, surprisingly promoted the formation of electrophysiologically silent synapses. Mechanistically, circRERE knockdown resulted in a preferential upregulation of synaptic mRNAs containing binding sites for miR-128-3p because of a reduced protective interaction between miR-128-3p and circRERE. Accordingly, overexpression of circRERE rescued exaggerated synapse formation upon circRERE knockdown in a miR-128-3p binding site-specific manner. Overall, our results uncover circRERE-mediated stabilization of miR-128-3p as a novel mechanism to restrict the formation of silent excitatory synaptic co-clusters and more generally implicate circRNA-dependent microRNA regulation in the control of synapse development and function.

## Introduction

Noncoding RNAs (ncRNAs) play increasingly appreciated gene-regulatory roles in the brain (Soutschek and Schratt, 2023) and can exist in both linear and circular forms. circular RNAs (circRNAs) are characterized by their stability and closed ends which are formed through a back-splicing event in which a covalent bond forms between a pre-mRNA 5’ splice donor and a 3’ splice acceptor site, leading to the formation of a unique backsplice junction sequence (BSJ) (Jeck et al., 2013) (Memczak et al., 2013). Recent work has further identified the dedicated mechanism by which circRNAs are exported to the cytoplasm dependent upon Ran-GTP/Exportin-2, underscoring the potential relevance of circRNA localisation for cell function (Ngo et al., 2024).

The circRNA transcriptome in the nervous system is particularly diverse and abundant (Gokool et al., 2019). circRNAs are highly expressed in developing neurons, often localized to the synapto-dendritic compartment and many are derived from genes with synaptic functions (Rybak-Wolf et al., 2014). Moreover, circRNAs are regulated in a neuronal activity-dependent manner (You et al., 2015), together suggesting that they could be involved in the control of synaptogenesis and plasticity. However, the function of the vast majority of the hundreds of conserved circRNAs expressed in mammalian neurons is still unexplored.

Arguably the most intensely studied neuronal circRNA is complementary determining region 1 antisense (Cdr1-as). Cdr1-as contains more than 70 binding sites for the neuronal microRNA (miRNA) miR-7 and acts as miR-7 sponge (Hansen et al., 2013), as part of a complex non-coding regulatory network (Kleaveland et al., 2018; Piwecka et al., 2017). The loss of Cdr1-has been shown to lead to miR-7 destabilization, thereby affecting excitatory synaptic transmission and schizophrenia-associated behaviour (Piwecka *et al*., 2017). Moreover, localised Cdr1-as activity is required for fear memory extinction (Zajaczkowski et al., 2023). In addition to Cdr1-as, a few additional circRNAs (e.g., circSatb1, circFat3, circGRIA1, circHomer1) have been implicated in the regulation of synapse development, plasticity and cognition (Gomes-Duarte et al., 2022; Seeler et al., 2023; Xu et al., 2020; Zimmerman et al., 2020).

Besides their role in regulating miRNA activity (Hansen *et al*., 2013), circRNAs have been shown to act through a variety of mechanisms including transcription and splicing regulation in the nucleus (Hollensen et al., 2020), RBP interaction (Ashwal-Fluss et al., 2014) and translation (Pamudurti et al., 2017).

Recently, neuronal circRNAs have been further implicated in neurological and neuropsychiatric diseases (Dong et al., 2023). For example, circIgfbp2 is significantly induced by traumatic brain injury (TBI), which is associated with several neurological and psychiatry disorders, and circIgfbp2 knockdown alleviated mitochondrial dysfunction and oxidative stress-induced synaptic dysfunction after TBI (Du et al., 2022). circHomer1 is downregulated in the prefrontal cortex of bipolar disorder and schizophrenia patients, and knockdown of circHomer1 in the mouse orbitofrontal cortex results in deficits in cognitive flexibility (Zimmerman *et al*., 2020). Furthermore, circRNAs are differentially expressed in pre- and post-symptomatic Alzheimer’s (Dube et al., 2019) and Huntington’s disease (Marfil-Marin et al., 2021).

CircRNAs are thus implicated in the regulation of synaptic plasticity and control of protein synthesis with relevance in the pathogenesis of neurodevelopmental and psychiatric disorders. However, the vast numbers, tissue specificity and splicing isoform complexity of circRNAs make it difficult to distinguish those with a functional role from a majority produced as presumably non-functional splice variants (Xu and Zhang, 2021). Therefore, we decided to investigate circRNA function in mammalian synaptogenesis in a more systematic manner, by first defining the circRNA landscape in the synapto-dendritic compartment, followed by RNA interference (RNAi)-mediated knockdown of synaptically enriched and activity-regulated circRNAs. This approach led to the identification of several candidate circRNAs for detailed mechanistic and functional investigation in follow-up experiments.

## Results

### Characterization of the circRNA landscape in the process compartment of primary rat hippocampal neurons

We reasoned that circRNAs which are highly abundant in neuronal processes (i.e., axons and dendrites) might represent strong candidates for circRNAs with functional relevance in synaptogenesis. To identify this neuronal circRNA population, we employed a previously characterized compartmentalised rat hippocampal neuron culture system, in which the transcriptome of neuronal somata and processes can be individually analysed after physical separation (Bicker et al., 2013).

By re-analysing our published ribo-depleted transcriptomic dataset (Colameo et al., 2021), followed by circRNA reconstruction of unique BSJs, we detected 1041 unique circRNAs, of which 907 circRNAs could be detected in the somata compartment and 1027 in processes (Figure 1A, Suppl. Table 1). Surprisingly, circRNA specific BSJ reads displayed a strong process compartment enrichment overall relative to the somata compartment when normalized to the total transcriptome (Figure 1B). This systematic process enrichment of circRNAs is also reflected upon normalization to the total reads of the circularRNA+linearRNA between the BSJ loci (Figure 1C), which in total showed a significant enrichment (FDR<0.05) in the process compartment for 234 circRNAs. This data suggests that neuronal circRNAs in general are biased towards synapto-dendritic localization, strongly implicating them in the regulation of local gene expression during synaptogenesis. However, circRNA dendritic enrichment likely occurs in a sequence-specific manner (Zajaczkowski *et al*., 2023), since several circRNAs were either not significantly enriched in the process compartment (e.g., Cdr1as) or even displayed strong soma enrichment (e.g. circSnap25) (Fig. 1D). We went on to validate selected process-enriched circRNAs using our previously established single-molecule fluorescent in situ hybridization (smFISH) protocol (Daswani et al., 2022). Thereby, we were able to confirm dendritic localization of the circRNAs circHomer1, circStau2 and circRMST in hippocampal pyramidal neurons (Figure 1E). The circularity of selected circRNAs was further validated by qPCR upon RNase R treatment (Suppl. Fig 1 A,B).

**Figure 1:**
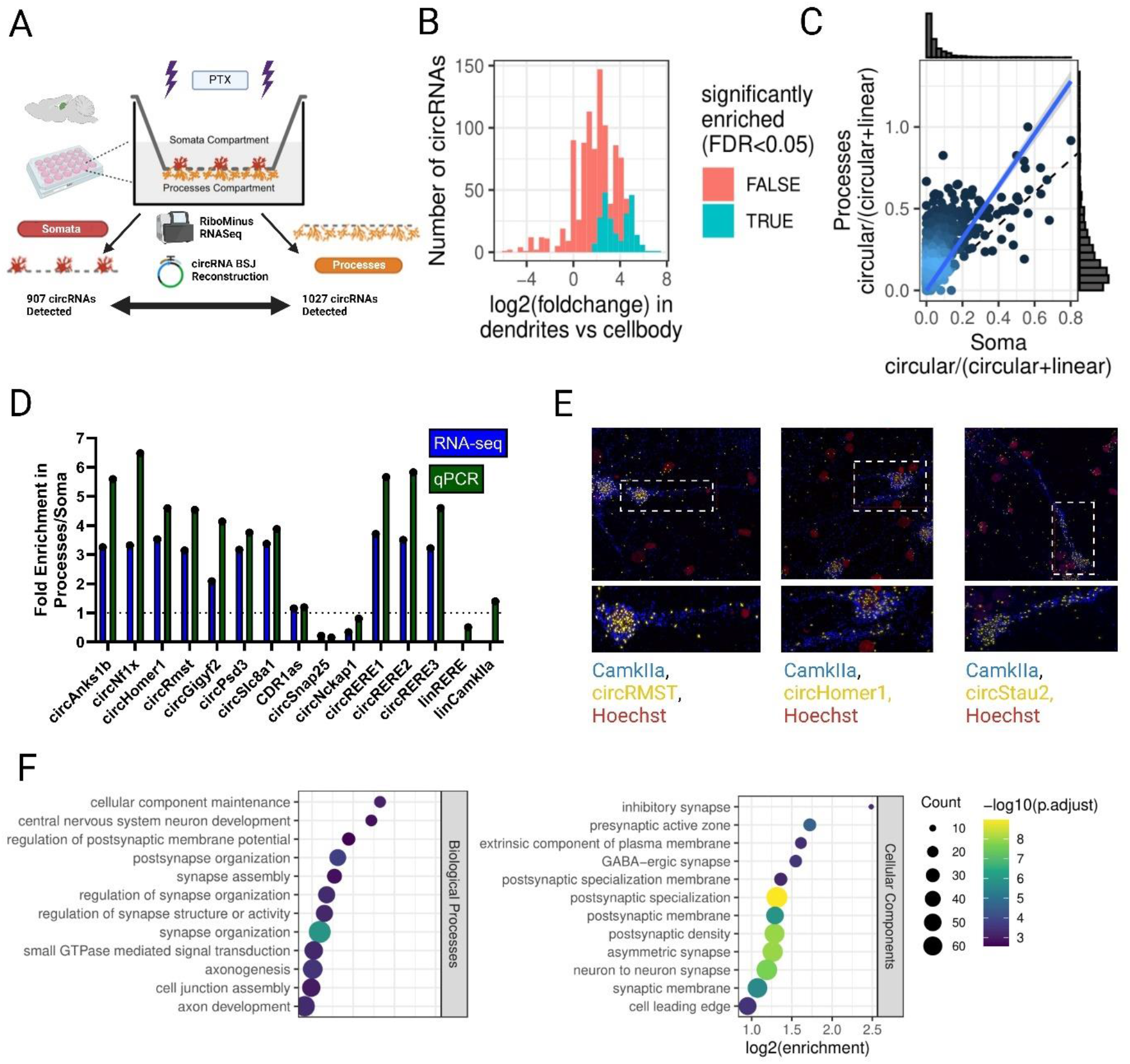
The circRNA landscape in processes of rat hippocampal neurons. A. Schematic of workflow for compartmentalized RNA extraction; circRNA BSJs were reconstructed from sequencing (DCC/Ciri) and quantified. B. Histogram of circRNA BSJ enrichment in the dendritic compartment normalized to total transcriptome, 4 replicates per group, 234/1041 circRNAs. C. circ/circ+lin ratios of circRNAs in processes and soma compartment, 4 replicates per group. Y-Axis illustrates significance of change (utilises a beta-binomial test (P Value <0.05) average read counts between somatic and process compartment, N=4). D. RT-qPCR with divergent primers for circRNAs from compartmentalized neuron cultures. circRNA enrichment in processes/soma is shown in comparison with average RNAseq result (n=1), Gapdh normalized. E. smFISH of selected circRNA candidates with probe designed against BSJ; Camk2a mRNA probe as neuronal marker. F. Gene set enrichment analysis for host genes of dendritically enriched circRNAs, using Biological Processes and Cellular Components ontologies.

Next, we decided to analyse the genetic origin of process-enriched circRNAs in further detail. Gene set enrichment analysis of circRNA host genes shows that process-enriched circRNAs are disproportionately often produced from synaptic genes, particularly those encoding post-synaptic proteins (Figure 1F). CircRNAs which are enriched in the process compartment are 4.2 times more likely to be on the SFARI autism risk gene set (65/234 circRNAs, p<2e-19) (Suppl. Table 2). Together, our bioinformatics results support a role for process-enriched circRNAs in synaptogenesis and synaptopathies, such as Autism Spectrum Disorder.

Most of the published examples of neuronal circRNAs act via miRNA association. However, miRNA sites in general do not appear to be overrepresented in circRNAs (Guo et al., 2014). In agreement with the latter result, circRNAs which are expressed in our neuron culture system on average do not contain more miRNA binding sites in comparison to mRNA 3’UTRs or whole transcripts (Suppl. Fig. 2 A-C). However, miRNA binding sites are slightly more abundant in dendritically enriched circRNAs compared to the total population (P<0.044) (Suppl. Fig. 2D). This effect is less pronounced when only considering 8-mer binding sites or when normalizing to the sequence length (Suppl. Fig. 2E, F). Higher abundance of miRNA sites appears to be restricted to a subset of circRNAs which overall contain a large number of binding sites (Suppl. Fig 2D, E), suggesting the presence of a specific circRNA sub-population with a high potential for miRNA regulation. Specific examples of process-enriched circRNAs containing many predicted miRNA binding sites are shown in Suppl. Fig. 2G,H.

Our ribo-depleted compartmentalized transcriptomic data (Colameo *et al*., 2021) further allowed us to interrogate activity-dependent regulation of neuronal circRNAs in the context of homeostatic plasticity (48hr Picrotoxin (Ptx) treatment at days in vitro (DIV) 18-20)). We found that chronic Ptx treatment changed the expression of a distinct subset of circRNAs (circular-to-linear ratio, FDR<0.1) (Suppl. Fig. 3 A/B) in a compartment-specific manner, e.g., circHomer1 (You *et al*., 2015) and several isoforms of the previously characterized circSlc8a1 (Hanan et al., 2020). However we observe distinctive expression of circRNA BSJs (Suppl. Fig. 3C) and the total exons of the circRNA tied to linear+circular counts, requiring further investigation to characterise their dynamic expression profiles (Suppl. Fig. 3D).

Taken together, processes of primary rat hippocampal neurons contain hundreds of circRNAs, some of which are regulated by neuronal activity and whose function in neuronal development is unexplored.

### RNAi screen reveals process-enriched circRNAs with a function in excitatory synaptic co-cluster and dendritic spine formation

Based on our results from RNA-sequencing, we decided to systematically characterize the function of process-enriched circRNAs in excitatory synaptogenesis. To narrow down the list of candidates to a number that can be handled in the context of a screen in primary neurons, we applied further selection criteria in addition to process-enrichment, namely absolute expression levels in processes (>10 counts in RNA-seq) and sequence conservation (orthologues in mouse and human).

This procedure led to the selection of 30 candidate circRNAs. Sometimes, this led to the inclusion of multiple isoforms from the same gene locus, e.g., in the case of circRERE, circANKS1B and Rmst (Fig. 2A). Additionally, the published miRNA regulatory circRNAs Cdr1-as (Hansen *et al*., 2013) and circHipk3 (Zheng et al., 2016), were added to the screen. siRNA pools (three independent siRNAs targeting the unique BSJ) were generated for each of the 32 circRNAs, as well as for two unrelated sequences for control purposes (siControl1, siControl2). Primary hippocampal neuronal cultures (DIV8) were transfected with GFP plasmid and siRNA pools followed by immunocytochemistry for synapsin-1/PSD-95 at DIV16, a time when excitatory synapse formation peaks in primary hippocampal neuron cultures. Subsequently, Synapsin-1/PSD-95 co-cluster density, which serves as a proxy for excitatory synapse density (Paradis et al., 2007), was automatically determined with a FIJI pipeline using GFP as a cell mask, as described previously (Inouye et al., 2022).

**Figure 2:**
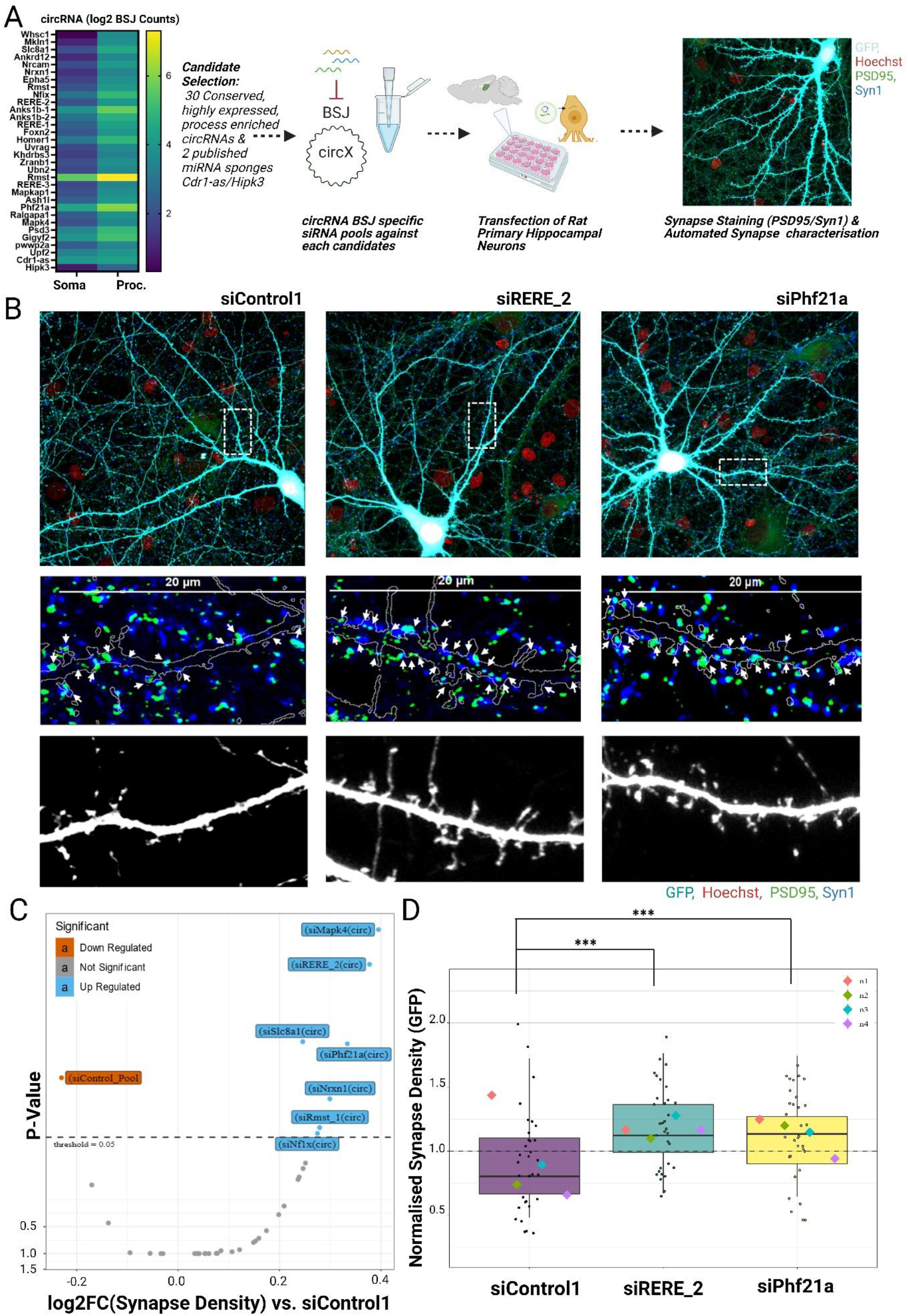
Several process-enriched circRNAs inhibit the formation of excitatory synaptic co-clusters and dendritic spines. A. Workflow of circRNA screen; siRNA pools designed against the BSJs of candidate circRNAs (30 process-enriched circRNAs plus Cdr1-as and Hipk3). Immunocytochemistry for pre (synapsin-1) and postsynaptic (PSD-95) marker was performed at DIV16, including GFP as a cell mask. B. Representative Images of selected circRNA candidates si-circPhf21a and si-circRERE2 with increased synaptic density upon knockdown compared to siControl; arrows depict synaptic co-clusters within the GFP mask. C. Volcano Plot showing fold-changes in PSD95/synapsin-1 co-cluster density for circRNA siRNA pools compared to si-Control1, GLMM statistical modelling, 4 independent biological replicates, 8 cells per condition. D. Boxplot of selected circRNA candidates for increased synapse density with noted pairwise comparisons (emmeans/GLMM).

In total, 7/32 siRNA pools against specific circRNAs resulted in a significant increase in synapse density, suggesting that these circRNAs act as endogenous repressors of synapse formation and/or maintenance (Fig. 2B-D; Suppl. Fig. 4A). Importantly, increased synapse density for these candidates was observed when using independent siControls for comparison (Suppl. Table 3), illustrating high robustness of our findings. To obtain independent support for a role of circRNAs in excitatory synapse development, we performed a secondary screen using as a read out dendritic spines, the major postsynaptic sites of excitatory synaptic contact. In this analysis, 11/32 candidates showed increased dendritic spine density when compared to siControl conditions (Suppl. Fig. 4B, C; Suppl. Table 3), with siRNA pools against circRERE2 and circPhf21a being the only conditions displaying an increase in both spine density and synapse co-cluster density. When analysing dendritic spine volume (Suppl. Fig. 4D, E; Suppl. Table 3), the only circRNA which resulted in a phenotype was Cdr1-as, whose knockdown significantly reduced dendritic spine size. In contrast, transfection of siRNA pools against circRERE2 and circPhf21a had no significant effect on dendritic spine volume, suggesting a specific effect on excitatory synapse formation. In conclusion, results from our siRNA screen suggest that several process-enriched neuronal circRNAs control different aspects of excitatory synapse development. Among those, circRERE and circPhf21a represent particularly strong candidates since they affect both the formation of excitatory synaptic co-clusters and dendritic spines.

circPhf21a has been previously identified as a highly expressed circRNA (2^nd^ most highly expressed circRNA in the rat brain, circAtlas 3.0 (Wu et al., 2024)). It is spliced from the 5’ UTR of Phf21a pre-mRNA and is enriched in the brain and in synaptoneurosomes (Rybak-Wolf *et al*., 2014). In contrast, much less is known regarding a neuronal function of circRERE. The RERE gene locus produces a wide variety of circular RNA isoforms which together outnumber the mRNA transcripts produced by the gene (Wu *et al*., 2024). Intriguingly, two of these circRERE isoforms containing different BSJs (circRERE1, circRERE2) are highly process-enriched in rat hippocampal neurons (Fig. 2A). Unlike circRERE2, circRERE1 KD was not statistically significant in our screen but nevertheless, circRERE-1 isoform knockdown similarly resulted in an increase in synapse (P<0.14) and spine density (P<0.19) (Fig. 2C, D, Supp Fig. 4B,C). We therefore decided to focus on circRERE for our further mechanistic studies.

### circular RERE isoforms suppress the formation, but promote the functional maturation of excitatory synaptic co-clusters

The RERE locus produces multiple circRERE isoforms due to different back-splicing events (Fig. 3A). This feature is evolutionarily conserved, with illustrated comparisons from rat (this study), mouse (Zajaczkowski *et al*., 2023) and human circRERE (Soutschek et al., 2023) Three structurally similar isoforms (circRERE1, circRERE2 and circRERE3) which are back-spliced out of exons 5-10/11/12 of the RERE gene are the most expressed process enriched circRERE isoforms (Figure 3A, B). circAtlas3.0 data repository counts circRERE1 (circAtlas ID: rno-Rere_0005 or circRERE (5,6,7,8,9,10)) and circRERE2 (circAtlas ID: rno-Rere_0019 or circRERE (5,6,7,8,9,10,11) as the 13^th^ and 28^th^ most highly expressed circRNAs in rat brain, respectively (Wu *et al*., 2024). Transcriptomic analysis of all RERE related transcripts demonstrates an overall process enrichment, supporting circRERE isoforms to be the predominant expressed RNA type in neurons when compared to linear RERE mRNA (Suppl. Fig. 5A). circRERE 1, 2 and 3 isoforms were also shown to be circular by resistance to RNase R degradation, in contrast to linRERE (Suppl. Fig. 5B). Rolling circle amplification (Das et al., 2019) of circRERE2 utilising BSJ spanning divergent primers demonstrates its circularity and confirms the expected size of 691nt size (Suppl. Fig. 5C). circular and linear RERE isoforms are relatively uniformly expressed in different rat brain regions (Fig. 3C). Over the course of rat hippocampal neurons differentiation *in vitro*, both linear and circular RERE isoforms initially decline, but peak at times of synapse formation (15-25 DIV; Fig. 3D). Furthermore, circRERE and linear RERE mRNA share comparable expression patterns in human iPSC neurons (Suppl. Fig 5E) (Soutschek *et al*., 2023), with circRERE1 and circRERE2 being the dominant circular isoforms.

**Figure 3:**
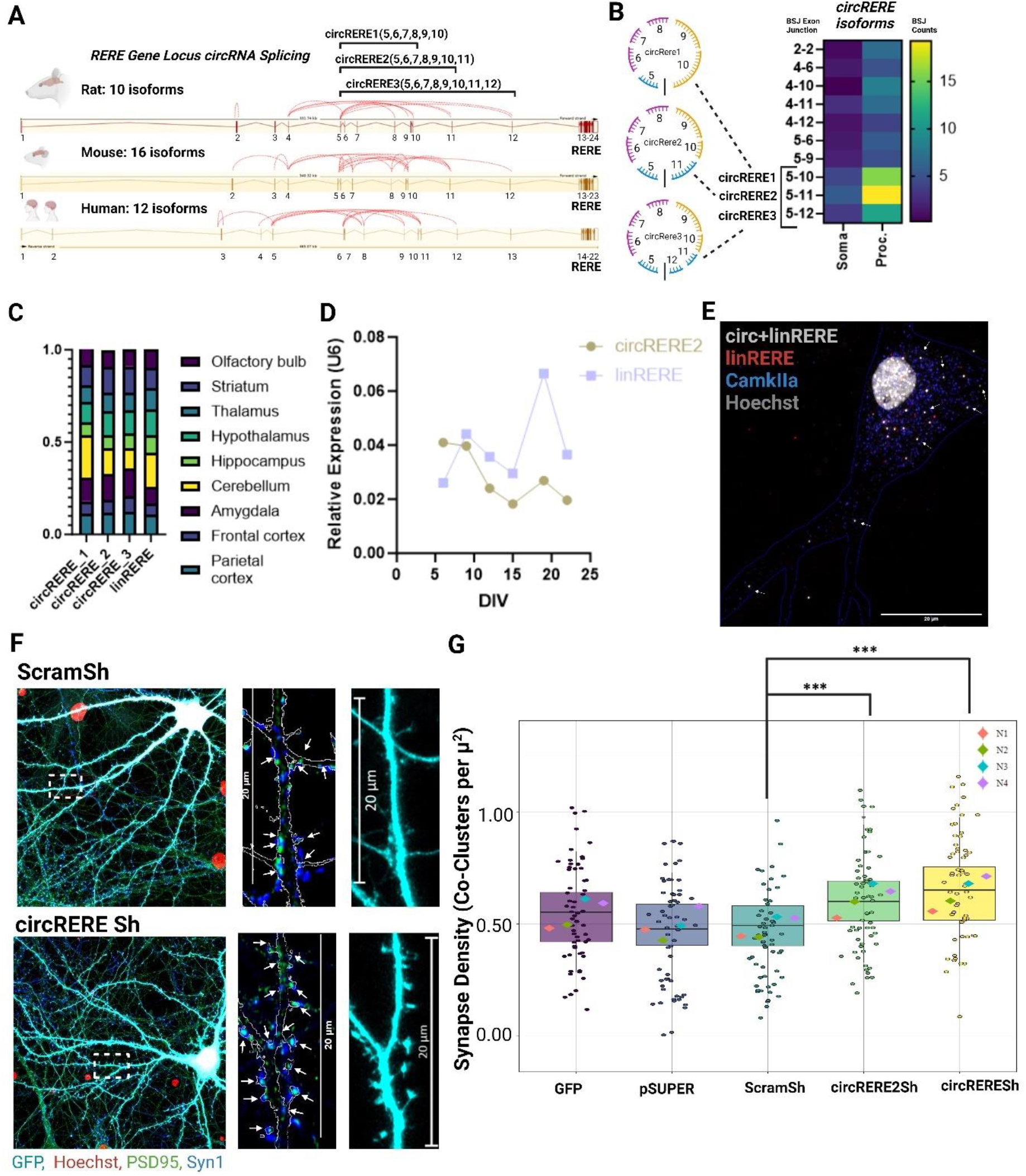
circRERE isoforms are abundant in neuronal processes and negatively regulate excitatory synapse co-clusters. A. Illustration of the conserved RERE gene locus and circRNA backsplicing between species, with emphasis on the major process-enriched isoforms circRERE1,2 and 3. B. circRERE isoform dendritic and soma expression heatmap comparing the unique BSJ counts, showing highest expression of circRERE1,2 and 3. C. circRERE isoforms and linRERE relative expression in adult rat brain (N=1). D. circRERE2 and linRERE expression timecourse in primary HC neurons (N=1), U6 Normalization. E. circRNA smFISH using probes directed against exons which are either specific for the linear RERE mRNA (exons 2-4, linRERE, red) or common to both circRERE and linear RERE mRNA (exons 5-10; circ+lin RERE, grey). Cytoplasmic circRERE puncta (grey only) are marked with arrows. F. Representative images from the pool of neurons analysed in (G) illustrating increased PSD95/Syn1 co-cluster density upon circREREsh. Arrows in blow-ups depict synaptic co-clusters within the GFP mask. G. Quantification of PSD95/Synapsin co-clusters in rat hippocampal neurons (DIV16) transfected with the indicated shRNAs or control plasmids. 15 Cells per condition per replicate, N-4 independent biological replicates. GLMM statistical modelling P<0.001 ***. Arrows in blow-ups depict synaptic co-clusters within the GFP mask.

We performed smFISH targeting the circRERE2 BSJ to specifically detect the subcellular localization of the most abundant RERE isoform in rat neurons (Suppl. Fig 5D). However, this approach only yielded a limited number of circRERE-2 positive puncta which were confined to the neuronal soma, likely due to the weak hybridisation of the FISH probe to the circRERE-2 BSJ. We therefore performed a dual smFISH with two different probes, one directed against exons 5-10 of the RERE gene (present in both the circular and linear isoforms (white) and one directed against exons 2-4 present only in linear RERE transcripts (red). From this, circRERE signal can be inferred by mutual exclusion. This analysis demonstrates circRERE localization not only in the nucleus and cytoplasm, but also in dendritic processes (Figure 3E; arrows).

Given the prominent expression of three circRERE isoforms with similar structural characteristics, we wondered whether there might be an additive effect on synaptogenesis by simultaneously knocking down circRERE isoforms 1-3 (referred to as circRERE sh from here on). The specificity and efficiency of the individual shRNA constructs targeting the different isoforms was validated by qPCR after transfection through nucleofection in rat cortical neurons (Suppl. Fig. 6A-F). circRERE2 knockdown alone resulted in a significant increase in synapse co-clusters compared to the ScramSh control (Fig. 3F, G), thereby confirming our results from the siRNA screen (Fig. 2). However, the knockdown of all 3 circRERE isoforms resulted in a more robust increase in dendritic synapse density compared to the circRERE2 shRNA (Figure 3G), indicating that circRERE isoforms 1 and 3 contribute to the repression of excitatory synapse formation, albeit to a lesser extent compared to circRERE2. The effect was specific for synapse formation since siRNA-mediated knockdown of neither of the circRNA isoforms 1-3 had a significant impact on dendritogenesis based on Sholl analysis (Suppl. Fig. 7 A-E). In contrast, linRERE knockdown resulted in widespread neuronal death of transfected cells, preventing synaptic characterisation, and suggesting that the observed increase in synapse co-clusters is not an artifact due to erroneous knockdown of the linear RERE mRNA.

Next, we explored whether increased synapse co-cluster density upon circRERE knockdown was accompanied by alterations in synaptic transmission using whole cell patch-clamp electrophysiological recordings. circREREsh transfected rat hippocampal neurons displayed a robust (49%) decrease in mEPSC frequency, but no significant change in mEPSC amplitude (Fig. 4A-E). This result implies that the vast majority of excess synaptic co-clusters formed upon circRERE knockdown might be functionally inactive (“silent”). We therefore decided to characterize excitatory synapses in these neurons in further detail. When plotting the mEPSC decay times, which depend on the subunit composition of AMPA-type glutamate receptors (AMPA-Rs) (Jonas, 2000), we observed a significant decrease in circREREsh compared to ScramSh transfected neurons (Fig. 4F,G), consistent with reduced GluA1 expression at circRERE knockdown synapses (Lu et al., 2009). Notably, genetic deletion of GluA1 leads to the formation of non-functional, “silent” synapses (Selcher et al., 2012). To assess GluA1 expression in circRERESh knockdown neurons more directly, we performed immunocytochemistry for surface-expressed GluA1 subunit (Fig. 4H,I). Transfection of both circRERE2sh and circREREsh resulted in a significant decrease in GluA1 expressed on the cell surface, with a more pronounced reduction (20%) upon circREREsh transfection (Fig. 4I). Similar to the results from synapse co-cluster density analysis (Fig. 3F,G), this again suggests a combined effect of multiple circRERE isoforms on the promotion of surface GluA1 expression. Together our results indicate a dual role of circRERE isoforms in excitatory synapse development by controlling both their formation and functional maturation, although in opposite directions.

**Figure 4:**
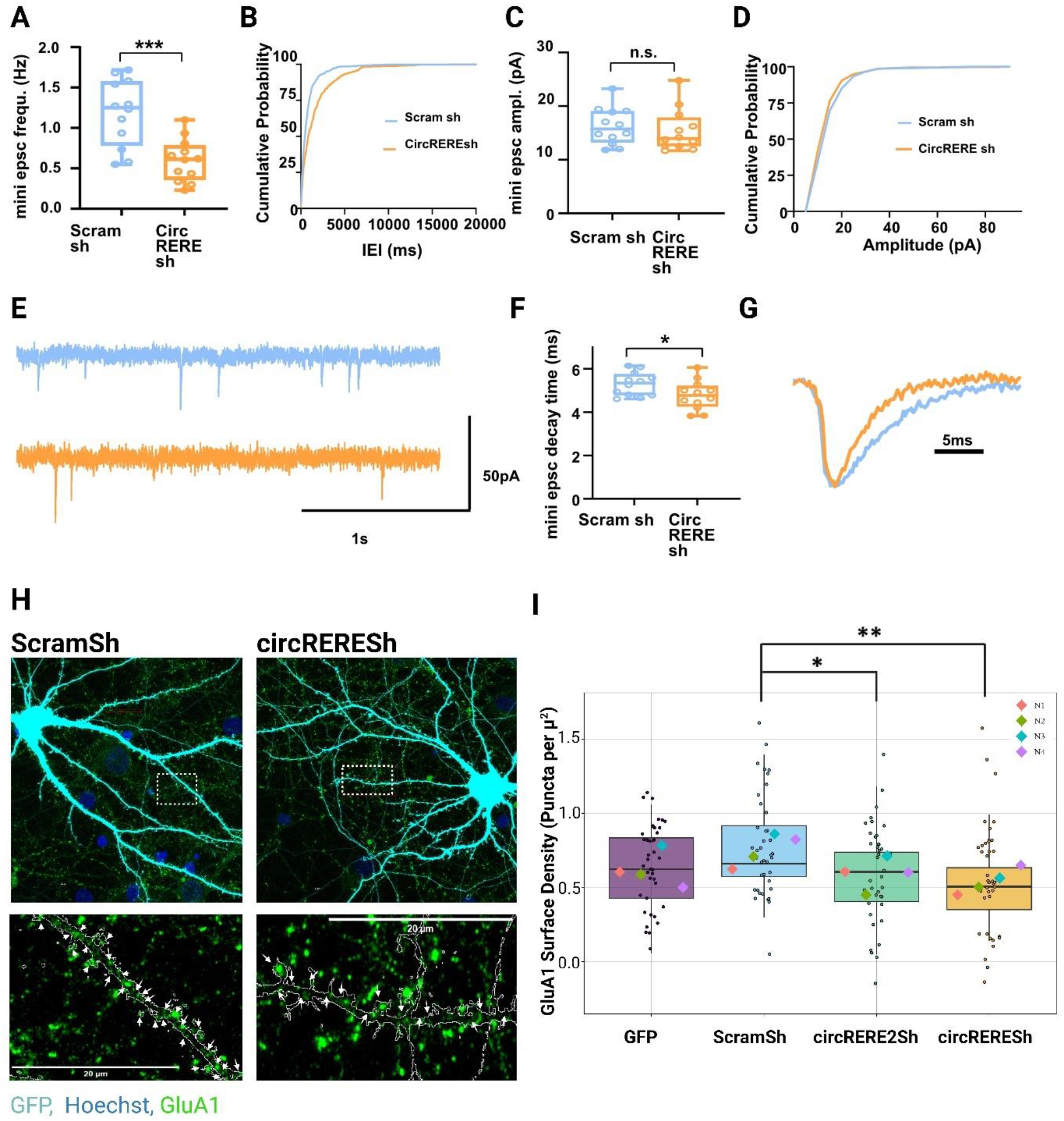
circRERE knockdown in hippocampal neurons leads to reduced excitatory synaptic transmission and GluA1 surface expression. Electrophysiological characterization of circRERE kd neurons. A/B. mEPSC frequency, C/D. mEPSC amplitudes. E. Representative traces supporting mEPSC frequency decrease with circREREsh compared to ScramSh. F/G. mEPSC decay time in circRERE shRNA (orange bars) or shRNA control (blue bars), N=12 neurons per condition. Unpaired T-test, P<0.05 *, P<0.001 ***. H. Representative images from the pool of neurons analysed in (I) illustrating reduced GluA1 surface puncta density upon circREREsh. Arrows depict GluA1 surface clusters within the GFP mask. I. Quantification of GluA1 surface expression in rat hippocampal neurons (DIV16) transfected with the indicated shRNAs or control plasmids. N4 independent biological replicates, 10 cells per condition per experiment, GLMM statistical modelling. P<0.05 *, P<0.01 **.

### circRERE suppresses excitatory synapse formation by stabilizing miR-128-3p

circRNAs are able to affect gene expression by a variety of different mechanisms (Kristensen et al., 2019) (Liu and Chen, 2022). Therefore, we reasoned that we might obtain first insight into the mechanism of circRERE function by assessing the protein-coding transcriptome in neurons upon circRERE knockdown. Primary rat cortical neurons were electroporated with the circRERE2sh or two control constructs (empty vector (pSUPER), ScramSh), followed by total RNA extraction and polyA-RNA sequencing. Bioinformatic analysis of the RNA-seq data revealed a total of 869 genes differentially expressed (DEGs) between the circRERE2sh and control conditions (Fig. 5A). 374 were upregulated, 495 were downregulated.

**Figure 5:**
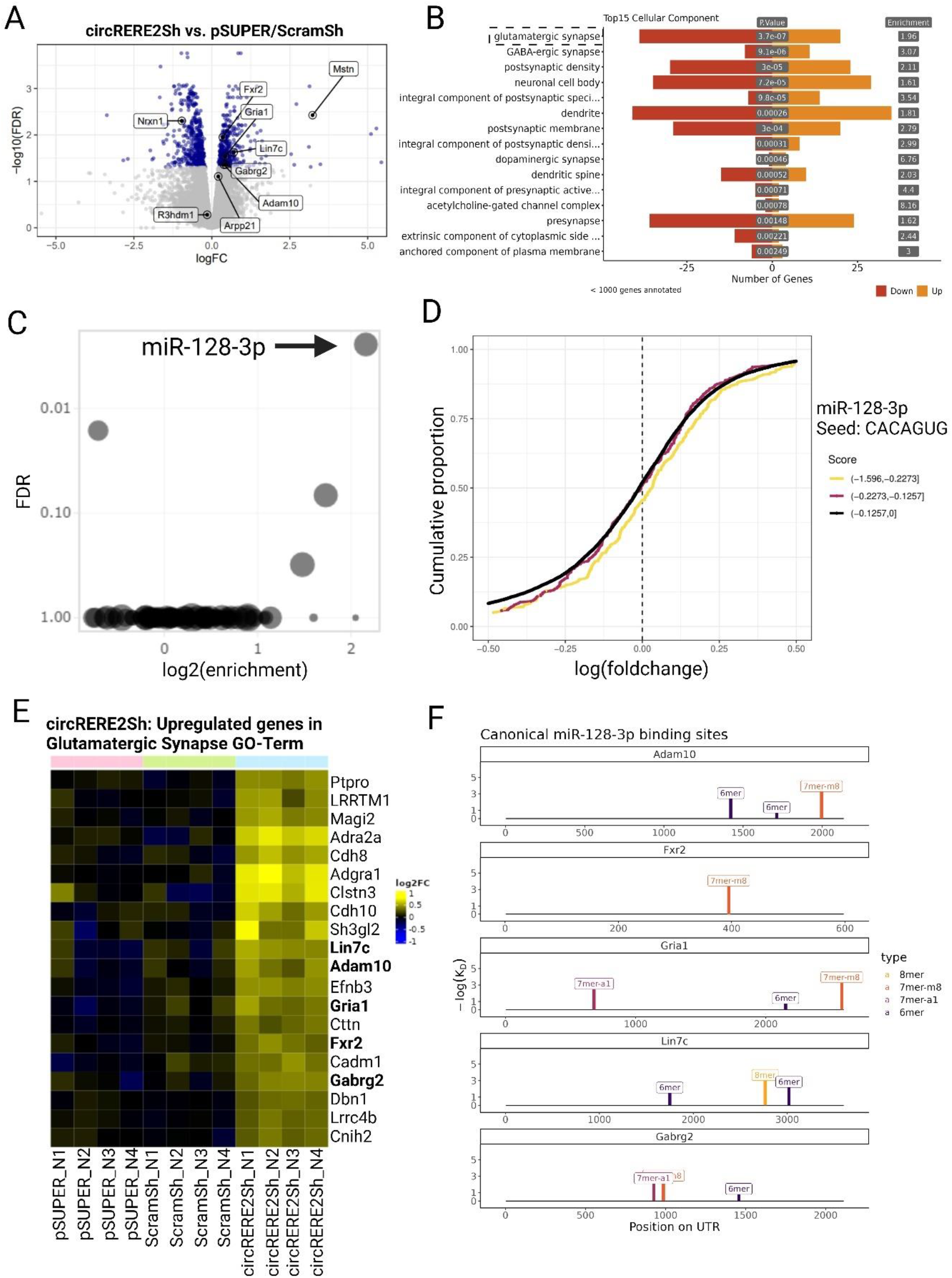
circRERE2 kd results in a preferential upregulation of miR-128-3p targets involved in excitatory synapse function. A. Volcano plot showing differentially expressed genes (DEGs) between circRERE2sh and control pSUPER/ScramSh electroporated rat cortical neurons (DIV5). Statistical analysis was performed with Genewise Negative Binomial Generalized Linear Models. B. Barplot showing number of genes in each significant GO term from cellular compartment ontology (ordered by elim); orange bars = upregulated genes, red bars = downregulated genes. C. EnrichMiR analysis of miRNA binding sites enriched in DEGs shown in A. D. Cumulative distribution (CD)-plot of log-fold changes from DEGs shown in A., containing either low (black line), medium (red line) or high (yellow line) affinity miR-128.3p binding sites. Right shift of the curves indicates on average higher expression of miR-128-3p containing transcripts in circRERE knockdown neurons. E. Heatmap of DEGs shown in A (FDR<0.05, upregulated only) comprised within the glutamatergic Synapse Go-Term. F. Illustration of miR-128-3p binding sites (separated by site type) within the 3’ UTR of the indicated upregulated DEGs.

GO Term analysis of the DEGs was performed to delineate biological pathways regulated downstream of circRERE (Fig. 5B, Suppl. Fig. 8 A,B). Strikingly, the top 15 GO terms in the category “cellular component” are mostly related to the synapse, with the top term being “glutamatergic synapse” (Fig. 5B). This was also supported by GSEA analysis with (GOCC_Synapse) as the stand-out gene set (Suppl. Fig. 8C). Together, these results are consistent with our results from the synaptic co-cluster analysis (Fig. 3E,F) and electrophysiology (Fig. 4A-G) and further support a role for circRERE in excitatory synapse development.

We were interested in how circRERE controls the expression of synaptic genes. Previously, several circRNAs have been shown to act as “miRNA sponges” by directly associating with specific miRNAs, e.g. Cdr1-as (Hansen *et al*., 2013) and circHipk3 (Zheng *et al*., 2016). We therefore used our previously developed bioinformatic webtool enrichmiR (Soutschek et al., 2022a) to assess whether specific miRNA binding sites are over-represented in DEGs over background. This would indicate altered activity of the corresponding miRNA in response to circRERE knockdown (kd). EnrichmiR analysis revealed that particularly transcripts containing binding sites for the neuronal miRNA miR-128-3p are significantly overrepresented in upregulated genes upon circRERE2 kd (Figure 5C). When plotting the expression fold-change for all genes as a function of the presence of miR-128-3p binding sites of different affinity (CD plot, Fig. 5D), we noticed that transcripts harboring particularly strong miR-128-3p binding sites (i.e. 8-mers) were on average more strongly upregulated in the circRERE kd neurons compared to those containing either weak or no sites. Thus, circRERE kd presumably leads to a lower miR-128-3p repressive activity, which is consistent with a stabilizing effect on miR-128-3p that is mediated by circRERE. As circRERE2kd presumably results in a reduction in miR-128-3p levels/activity, we focused on upregulated genes in the synapse GO term (n=20; Figure 5E). Among those, five (Adam10, Fxr2, Gria1, Lin7C and Gabrg2) contain canonical, high-affinity miR-128-3p binding sites in their 3’ UTR (Figure 5F). Thus, upregulation of this group of synaptic genes upon perturbation of a protective circRERE/miR-128-3p interaction might underlie the observed synaptic phenotypes.

Previously, knockout of the neuronal Cdr1as was shown to lead to a de-stabilization of its binding partner miR-7 (Piwecka *et al*., 2017). To investigate the impact of circRERE knockdown on the expression of mature miRNAs, we subjected the same samples used for polyA RNA-seq to small-RNA-seq. Differential expression analysis revealed that the expression of miR-128-3p (rno-miR-128-2) (Fig. 6A) was indeed highly significantly reduced. Based on polyA-RNAseq, transcripts originating from both of the miR-128-3p host genes (Arpp21, R3hdm1) are not significantly downregulated in circRERE knockdown neurons, arguing against an effect on miR-128-3p transcription or processing. In fact, the host of 128-2-3p, Arpp21, is itself regulated by 128-3p via a negative feedback process (Rehfeld et al., 2018). Accordingly, an upregulation of Arpp21 (FDR<0.0674) can be observed upon circRERE knockdown. R3hdm1, the host of miR-128-1-3p, is unaffected (Suppl. Fig. 8D). Together, our results are consistent with a role for circRERE in the stabilization of the mature miR-128-3p.

**Figure 6:**
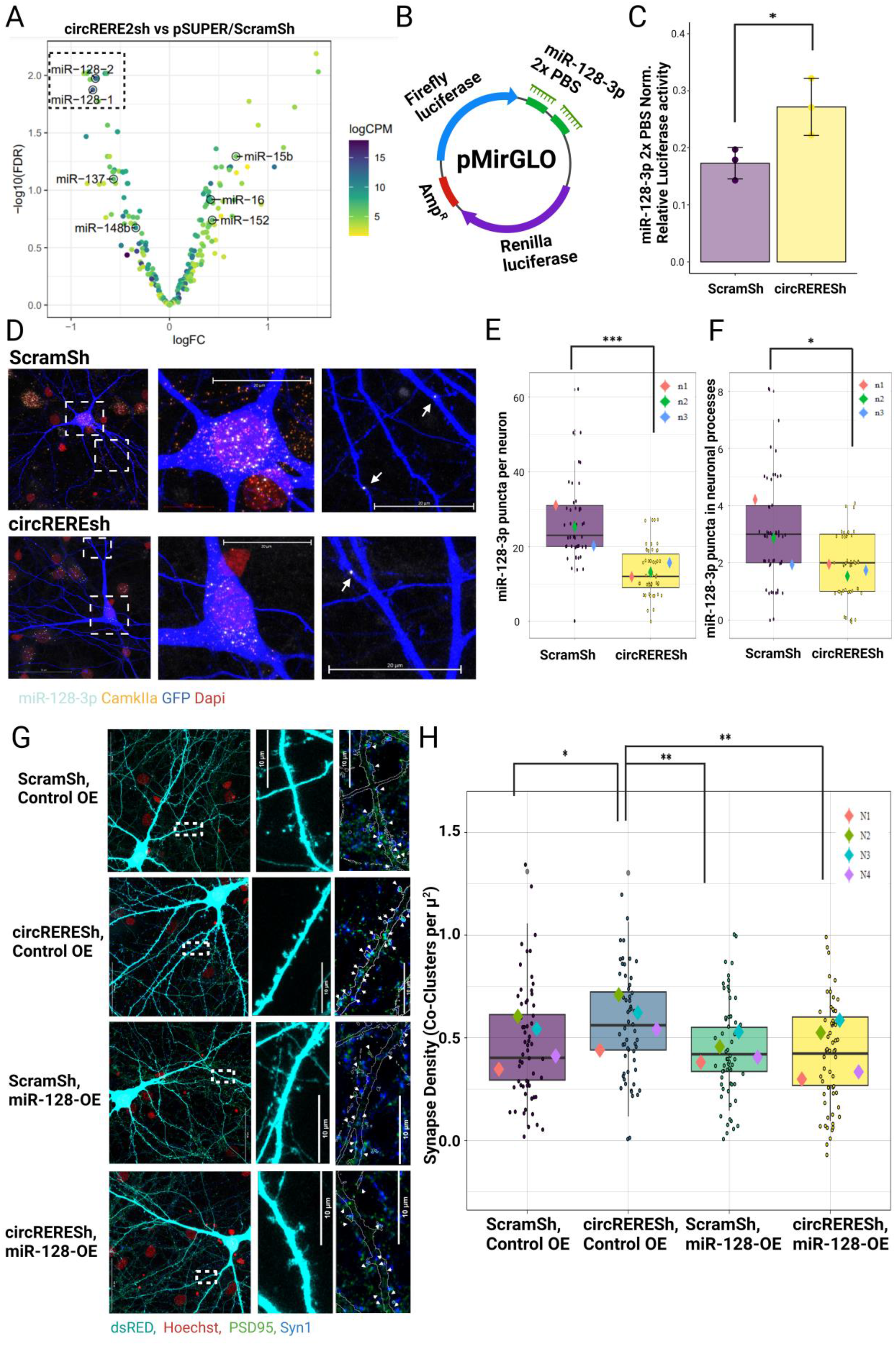
circREREkd results in reduced miR-128-3p expression levels and activity. A. Volcano plot showing differentially expressed miRNAs between circRERE2sh and control shRNA. Statistical analysis was performed with Genewise Negative Binomial Generalized Linear Models. In case of multiple miRNA isoforms, only one was labeled. B. Illustration of 2X Perfect binding site Dual-luciferase construct sensitive to miR-128-3p activity (miR-128-3p 2x PBS). C. Relative luciferase activity in neurons transfected with miR-128-3p 2x PBS together with the indicated shRNA constructs. N=3, unpaired t-test, P<0.05. D. Representative images from double smFISH experiment in ScramSh (upper row) or circREREsh (lower row) transfected rat hippocampal neurons using probes directed against miR-128-3p and Camk2a. Arrows point to miR-128-3p positive puncta in dendrites. E. Quantification of miR-128-3p puncta in somata or dendrites (F.) of neurons transfected as in C, n=3 GLMM, 15-20 cells per condition per replicate. P<0.05 *, P<0.001 ***. Each datapoint represents the number of puncta detected in an individual neuron. G. Representative images from the pool of neurons analysed in (H) depicting rescue of PSD95/Syn1 co-cluster density in circRERE KD neurons upon miR-128-3p overexpression. Arrows in blow-ups depict synaptic co-clusters within the GFP mask. H. Quantification of PSD95/Synapsin co-clusters in rat hippocampal neurons (DIV16) transfected with the indicated shRNAs and overexpression (OE) plasmids.15 Cells per condition per replicate, N= 12-16 Cells per experiment per condition, 4 independent experiments, GLMM statistical modelling. P<0.05 *, P<0.01 **.

Next, we addressed more directly whether the observed reduction in miR-128-3p levels upon circRERE knockdown translated into reduced miR-128-3p repressive activity. Therefore, we performed luciferase assays with a 2X perfect miRNA binding site reporter in the context of circREREsh (Fig. 6B,C). circRERE kd significantly increased reporter gene activity by 39% compared to controls, confirming that reductions in miR-128-3p expression indeed impair miR-128-3p repressive activity.

To visualise the subcellular localisation of miR-128-3p in neurons and to validate reduced miR-128-3p expression upon circREREsh, miRNA smFISH was performed (Fig. 6D-F). circREREsh resulted in an average 47% decrease in miR-128-3p positive puncta in hippocampal pyramidal neurons (identified by Camk2a positive puncta), highly consistent with our results from small RNA seq (Fig. 6A). Overall, the magnitude of reduction was similar between the somatic and dendritic compartment, although the vast majority of puncta (∼90%) were detected within the soma (Fig. 6F). This result agrees with previous literature, which did not report dendritic or synaptic enrichment of miR-128-3p in various preparations e.g. (Siegel et al., 2009). We note however that absolute quantification of miR-128-3p by qPCR in primary hippocampal neurons led to an estimate of about ∼4000 copies of miR-128-3p per neuron (de la Mata et al., 2015), which suggests that miRNA FISH is capturing only a subset of the total population (Fig. 6D-F). Thus, miR-128-3p is presumably present at sufficient quantity in dendrites to control the local expression of synaptic target mRNAs.

Given the pronounced effect of circRERE knockdown on miR-128-3p expression, we asked whether restoring miR-128-3p levels was sufficient to rescue the synapse phenotype caused by circRERE knockdown. Towards this end, we also utilised a previously published plasmid for miR-128-3p overexpression (dsRED-miR-128-3p, from here on called miR-128 OE) (Rehfeld *et al*., 2018). Using the 2x miR-128-3p PBS luciferase assay, we validated the expression of functional miR-128-3p by miR-128 OE (Suppl. Fig. 9A). Moreover, when we co-express miR-128-3p OE together with circREREsh (Fig. 6G,H), excessive synapse co-cluster density caused by circRERE knockdown alone returned to baseline levels, demonstrating efficient rescue. Interestingly, miR-128OE alone did not further decrease synapse density, possibly due to its high expression already at basal levels.

Together, our results strongly support the hypothesis that the synaptic phenotype caused by circRERE knockdown is mediated by a decrease in miR-128-3p levels.

### circRERE function in excitatory synapse formation is dependent on the presence of intact miR-128-3p binding sites

We wondered whether circRERE stabilizes miR-128-3p levels via a direct interaction, i.e. protecting it from destruction via endogenous decay pathways. Using ScanmiR, a bioinformatic tool developed in our lab that identifies predicted miRNA binding sites within transcripts (Soutschek et al., 2022b), we detected two putative high-affinity miR-128-3p binding sites within exon 9 of circRERE (Fig. 7A). Importantly, these sites are present in all of the highly expressed circRERE isoforms 1-3. In addition, an AGO HITS-CLIP peak is present on RERE exon 9 between these sites (Chi et al., 2009), further supporting a direct interaction between circRERE and miRNAs. To investigate the functional relevance of these two miR-128-3p binding sites, we generated circRERE overexpression constructs utilising the Wilusz lab’s Zkscan MCS OE vector (Kramer et al., 2015), containing RERE exons 5-12 (corresponding to circRERE3 with a different BSJ sequence), either containing wild-type (WT) or mutant (MUT) versions of the 7mer and 8mer miR-128-3p binding sites detected on exon 9 (Figure 7A). The efficacy and specificity of circRERE WT and circRERE 128-MUT construct overexpression was validated following electroporation of primary cortical neurons by RT-qPCR (Suppl. Fig. 9B). The empty Zkscan MCS OE Vector was also utilised as an additional control which does not generate circRERE. Endogenous circRERE 1, 2 and circRERE3 were detected using BSJ spanning primers which do not efficiently bind to the overexpressed circRERE due to its unique BSJ. Accordingly, endogenous circRERE1-3 detection were mostly unaffected by circRERE overexpression. More importantly, by utilising primers flanking exons 5-12 (circRERE-WT), we detected efficient overexpression of circRERE by both the circRERE-WT and circRERE 128-Mut constructs, which occurred to a similar extent.

**Figure 7:**
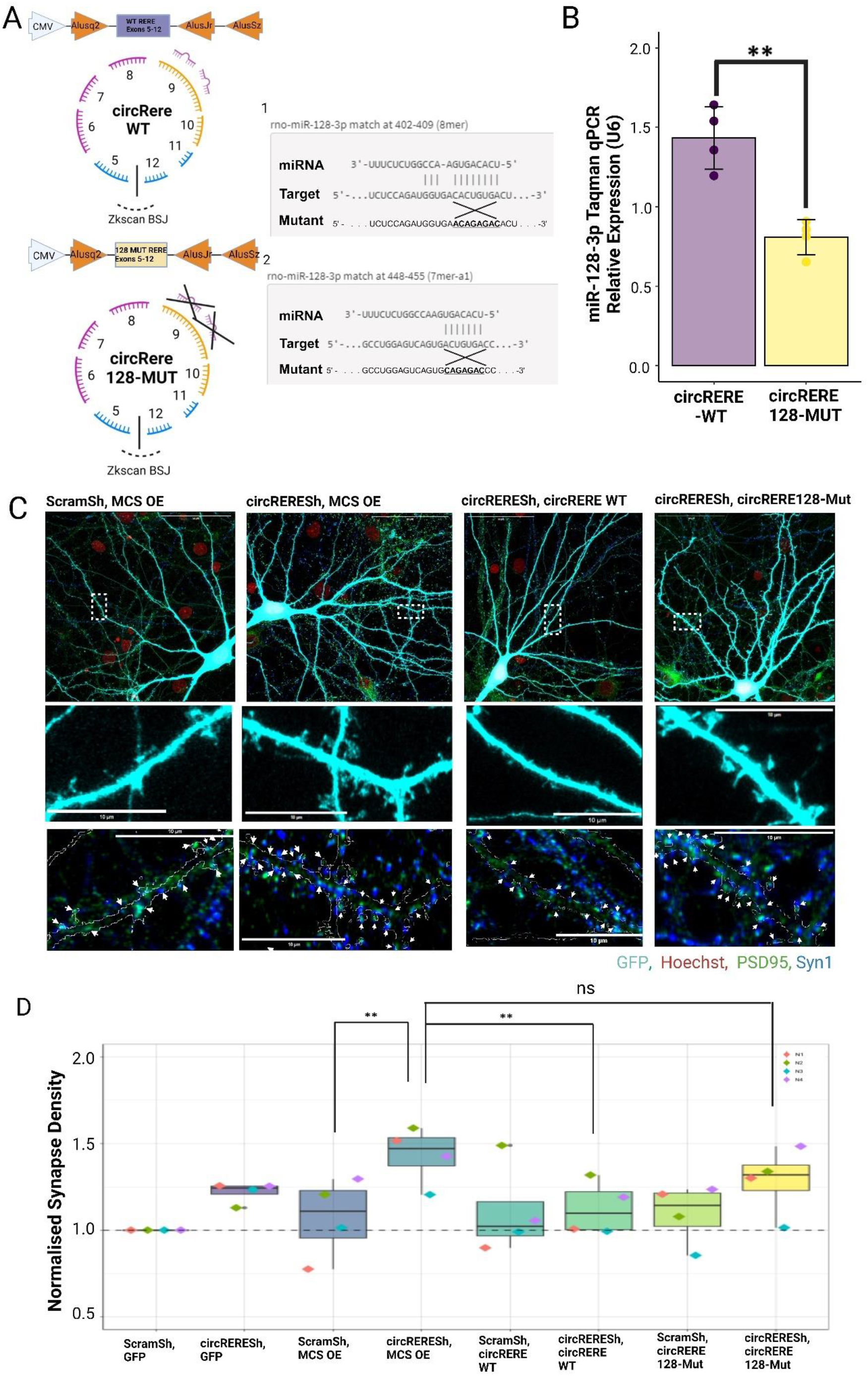
circRERE function in excitatory synapse formation is dependent on the presence of intact miR-128-3p binding sites. A. Schematic of the circRERE Wt and 128-Mut OE constructs. The two miR-128-3p binding sites within exon 9 of circRERE are shown in their wild-type and mutant configuration. B. miR-128-3p expression levels determined by Taqman miRNA qPCR in primary cortical neurons (DIV5) electroporated with either circRERE WT or 128-Mut OE constructs. U6 snRNA was used for normalization. Unpaired t-test P<0.01 **. C. Representative images from the pool of neurons analysed in (D) illustrating rescue of synaptic co-cluster density in circRERE KD neurons upon expression of circRERE WT, but not 128-Mut. Arrows in blow-ups depict synaptic co-clusters within the GFP mask. D. Quantification of PSD95/Synapsin co-clusters in rat hippocampal neurons (DIV16) transfected with the indicated shRNAs and circRNA expression plasmids. MCS: multiple cloning site. Pairwise comparisons from emmeans/GLLM. N4 biological replicates, 15 cells per condition, P<0.01 **; ns: not significant.

Next, we explored the effect of circRERE-WT overexpression on miR-128-3p expression by miRNA qPCR (Fig. 7B). circRERE-WT overexpression (with intact miR-128-3p binding sites) led to a highly significant, 48% increase in miR-128-3p levels compared to circRERE 128-Mut (containing mutated miR-128-3p binding sites) (Fig. 7B). This supports the hypothesis of a protective interaction between circRERE and miR-128-3p in a manner dependent upon the presence of the predicted two miR-128-3p miRNA binding sites.

Having shown the differential effects exerted by the circRERE -WT and 128-MUT constructs on miR-128-3p expression, we further tested their ability to rescue excessive synapse co-cluster density caused by circRERE knockdown. Importantly, both constructs are resistant to the circRERE shRNA due to their unique BSJ. We expected that an efficient rescue should only be achieved with the circRERE WT-overexpression, since the presence of miR-128-3p binding sites within circRERE is necessary for miR-128-3p stabilization. In agreement with this assumption, we observed a rescue of synapse co-cluster density with circRERE-WT, but not circRERE 128-mut, when co-expressed with circREREsh (Fig. 7C,D).

In conclusion, these results tie the presence miR-128-3p binding sites within circRERE to its effects on excitatory synapse formation. This strongly supports a model whereby circRERE protects mir-128-3p from degradation via direct interaction at baseline, thereby stabilizing neuronal mir-128-3p levels and restricting synapse formation via the downregulation of critical miR-128-3p synaptic target mRNAs at the peak of excitatory synaptogenesis *in vitro* (Fig. 8).

**Figure 8:**
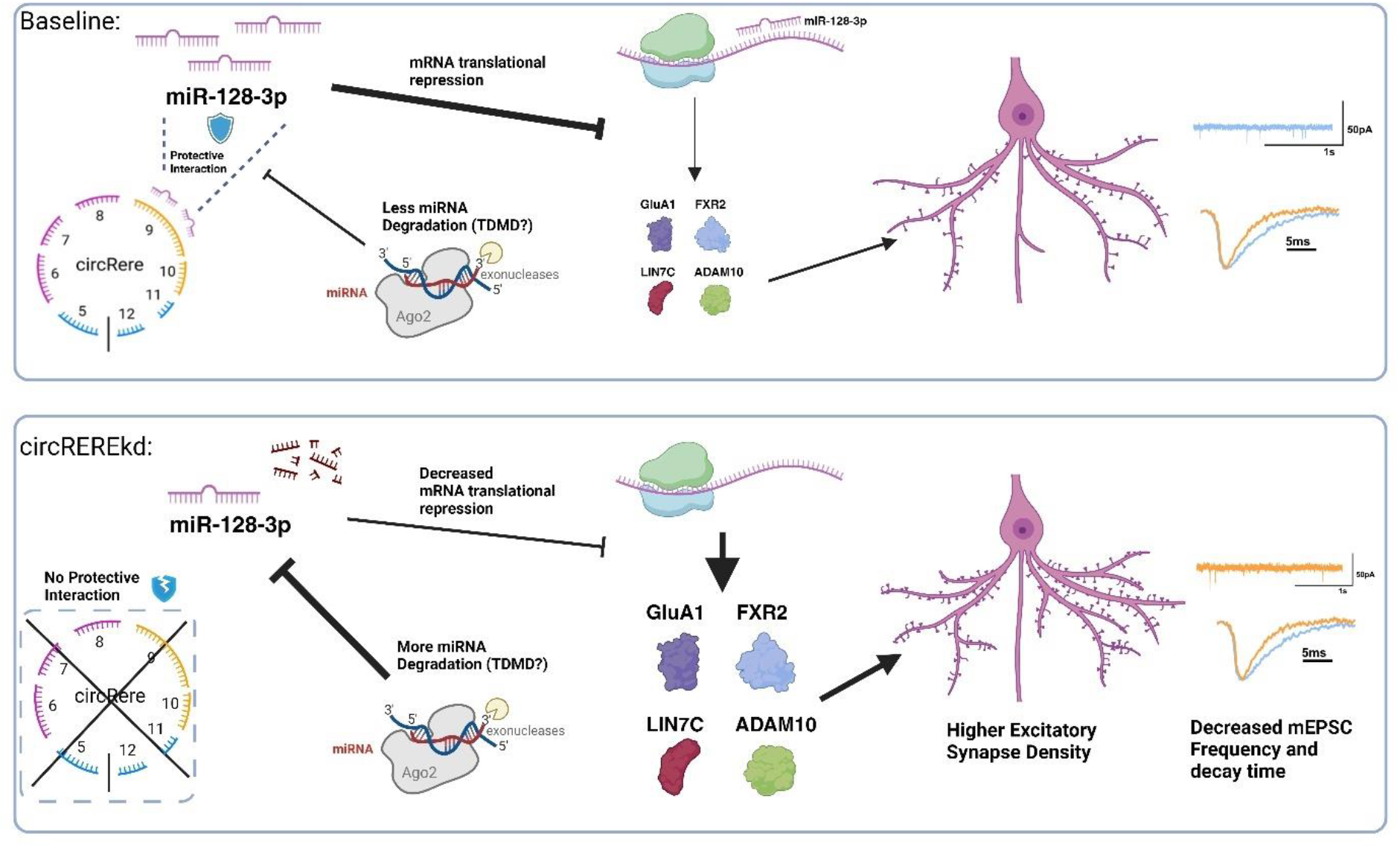
Proposed model of circRERE function. Under basal conditions, circRERE maintains a dynamic protective interaction with miR-128-3p, preventing a reduction in the active miR-128-3p pool, e.g., mediated by the target dependent microRNA degradation (TDMD) pathway. High miR-128-3p activity in turn is necessary for efficient translational repression of synaptic genes involved in synapse formation and/or maturation. Upon circREREkd, this protective interaction is lost, leading to enhanced miR-128-3p degradation, and reduced translational repression of synaptic target genes, increase in synapse density and decreased mEPSC frequency.

## Discussion

In this study, we performed the first systematic characterization of neuronal circRNAs during excitatory synaptogenesis. By using compartmentalized primary rat hippocampal neuron model, we identified several circRNAs as inhibitors of excitatory synapse formation, and further characterized the molecular mechanism of one of these candidates, circRERE. Thus, our study demonstrates a significant contribution of circRNAs to the regulation of mammalian synaptogenesis and provide new insight into their mode of action, specifically regarding their regulation of miRNA activity. This will form the basis for follow-up studies on their (patho)physiological relevance using *in vivo* models.

### circRNAs are enriched in the synapto-dendritic compartment

For the identification of process-enriched circRNAs, we leveraged a previously described primary rat hippocampal neuron model which allows the physical separation of processes (axons, dendrites) from somata. Since this system also contains glial processes (Bicker *et al*., 2013), circRNAs detected by our RNA-seq approach could in principle also originate from glial cells. However, 113/234 processed enriched circRNAs were also previously detected in a mouse synaptoneurosome preparation (Rybak-Wolf *et al*., 2014) including 8 circRERE isoforms. This includes 31/32 screened circRNAs in our study (with the exception of Ankrd12). Moreover, we (Fig. 1E) and others (You *et al*., 2015) could validate dendritic localization of several candidates by smFISH. Therefore, the vast majority of the process-enriched circRNAs is likely of neuronal origin.

One of the surprising findings of our work is that a large fraction (22%) of neuronal circRNAs displays a high enrichment in the process compartment (Fig.1B). This implies the existence of specific circRNA transport mechanisms which remain to be determined and might, for example, involve post-transcriptional m6A modifications (Qin et al., 2022) or the interaction with RNA-binding proteins (RBP) (Dell’Orco et al., 2020). Alternatively, since circRNAs are inherently more stable than other RNA types, they could gradually accumulate over time in the synapto-dendritic compartment. We favor the first model, since it is more consistent with the observation that a small subset of neuronal circRNAs (e.g, circSnap25, Capns1, Ubr5, Nckap1) is actively depleted from processes, which would argue against passive diffusion. What might be the functional relevance of this compartment-specific circRNA enrichment? One can speculate that it might skew the stoichiometric ratio of circRNAs and its interaction partners (miRNA or RBP) in favor of circRNAs, thereby allowing for a more efficient functional interaction, i.e., in the context of local protein synthesis regulation at synapses.

### circRNAs act as negative regulators of excitatory synapse formation

To our knowledge, this study represents the first systematic screen interrogating circRNA function in post-mitotic neurons. While systematic functional screens for circRNAs are generally scarce, RNAi based (Liu et al., 2022) and CRISPR-Cas13 based screens (Li et al., 2021) have been primarily undertaken in cancer cells. Interestingly, our main hit, circRERE, has been identified as a cell essential circRNA that regulates ferroptosis in cancer cell lines (Liu *et al*., 2022). Intriguingly, our screen results show that circRNAs have an overall repressive impact on synapse formation, possibly preventing the premature and/or excessive formation of synapses during neuronal development. When interrogating the functional implications with electrophysiological recordings for the candidate circRERE, we surprisingly found that increased synapse co-cluster density upon circRERE knockdown was associated with reduced mEPSC frequency. This finding is most consistent with the idea that these newly formed synaptic co-clusters are non-functional due to deficits in postsynaptic GluA1 expression (“silent synapses”). However, at this point we cannot exclude additional or sole deficits in presynaptic release. A more in-depth electrophysiological characterization is required to distinguish between these possibilities. In any case, this suggests a dual function of circRERE, working both as a repressor of synapse formation, but also as a promoter of synapse maturation. We speculate that such a function could be important to ensure the homeostasis of neural circuit activity during development.

Besides the circRNAs described here, only a few additional circRNAs have previously been shown to be involved in the regulation of synapse development. Cdr1-as is arguably the most characterized circRNA and has been shown to regulate the activity of miR-7 in an activity-dependent manner (Jara et al., 2023). Cdr1-as knockout in mice leads to dysfunction in excitatory synaptic transmission with an increase in spontaneous excitatory postsynaptic currents (Piwecka *et al*., 2017) congruent with our screen, where we picked up Cdr1-as as the top regulator of dendritic spine size and number (Suppl. Fig 4 B,C). Further examples include CircGria1, whose knockdown increases synaptogenesis in the macaque brain, presumably via upregulation of linear Gria1 mRNA from its host gene (Xu *et al*., 2020). On the other hand, the dendritically enriched circHomer1 was shown to regulate synaptic genes by interacting with the RBP HuD (Zimmerman *et al*., 2020), although the impact on synaptic transmission was not directly addressed in this study. Finally, circSatb1, which was not part of our screen, was shown to regulate dendritic spine morphology in hippocampal neurons (Gomes-Duarte *et al*., 2022), but the underlying mechanism and relevance for synaptic transmission were not addressed.

### circRERE function in synaptogenesis requires miR-128-3p

We provide multiple lines of evidence that circRERE function critically relies on the stabilization of the neuronal miRNA miR-128-3p (Fig. 8). miR-128-3p is abundantly expressed in rodent neurons (Epple et al., 2021) and has been repeatedly involved in the regulation of synapse development and function. In agreement with our observation of reduced miR-128-3p expression upon circRERE knockdown, miR-128-3p deficiency in D1 dopaminergic neurons results in increased dendritic spine number (Tan et al., 2013). However, miR-128-3p deficiency was further shown to lead to increased neuronal excitability in multiple studies, which culminates in an increased susceptibility for epileptic seizures (Tan *et al*., 2013) (Franzoni et al., 2015) (Rehfeld *et al*., 2018) (McSweeney et al., 2016) (Shvarts-Serebro et al., 2021) . This would suggest that the effects of circRERE on synapse formation and function are uncoupled and display differential dependency with regards to miR-128-3p. To address this, additional experiments on the impact of miR-128-3p on synapse physiology in the context of circRERE knockdown are required.

RNA-sequencing revealed interesting miR-128-3p targets which could mediate the positive effects of circRERE knockdown on excitatory synapse formation (Fig. 5). For example, FXR2 was shown to promote the activity-dependent local translation of PSD-95, a central component of the post-synaptic density (Fernandez et al., 2015). Gria1, encoding the AMPA-type glutamate receptor subunit GluA1, is a critical determinant of dendritic spine structure and excitatory synapse function (Diering and Huganir, 2018). Interestingly, despite increased Gria1 mRNA levels, we observed less GluA1 surface expression upon circRERE knockdown (Fig. 4H,I), which is in agreement with our results from electrophysiological recordings (Fig. 4A-G) and provides further support for the existence of different pathways controlled by circRERE during synapse formation and maturation. The matrix metalloprotease Adam10 has been shown to control the localization of GluA1 to synapses and to remodel dendritic spines through the cleavage of adhesion molecules, such as N-Cadherin (Gardoni et al., 2012) (Malinverno et al., 2010). It will be interesting to test which of these miR-128-3p targets, if any, is causally involved in circRERE-mediated effects on either synapse formation or functional maturation.

### circRERE acts via miR-128-3p stabilization

The reported molecular mechanisms of circRNA action are diverse, ranging from protein translation to protein binding and miRNA sponging, with the latter being arguably the most widely described. For most cases, a sponging interaction between circRNAs and miRNAs has been shown to reduce the activity of the corresponding miRNAs on their targets, e.g., circHipk3 and circSry (Zheng *et al*., 2016), (Hansen *et al*., 2013). However, the consequence of these interactions is not as straightforward as first envisioned (Fuchs Wightman et al., 2024). For example, Cdr1-as was originally conceived as a classical miRNA “sponge” (Hansen *et al*., 2013), but further work rather suggests a protective interaction between Cdr1-as and miR-7. According to this, Cdr1-as protects miR-7 from TDMD mediated degradation by the lncRNA Cyrano (Piwecka *et al*., 2017), (Kleaveland *et al*., 2018). A similar protective interaction has also been recently described for circCSNK1G3, which harbors strong TDMD-like sites for Mir-181b/d. However, despite increased miR-181 levels, circCSNK1G3 expression resulted in an overall decrease in target gene repression (Piras et al., 2022). In the case of circRERE, our data currently supports a stabilizing interaction between circRERE and miR-128-3p, similar to what was observed for Cdr1as and miR-7. Interestingly, despite its strong dendritic enrichment, circRERE knockdown affected miR-128-3p levels similarly in the dendritic and somatic compartment (Fig. 6F), likely due to the still abundant somatic expression of circRERE. Nevertheless, due to the different stoichiometry, the impact on miR-128-3p repressive activity might still vary between the different compartments. Future experiments which more directly address the local translation of miR-128-3p targets in the soma and dendrites, preferentially in the context of neuronal activity changes, should help to resolve this issue. Finally, one would postulate the existence of mRNAs which trigger TDMD of miR-128-3p in the absence of circRERE, analogous to Cyrano. Recently developed bioinformatics approaches should help to identify such triggers, which can subsequently be experimentally tested. On a broader perspective, it will be interesting to know how widespread this mechanism is, i.e. by studying additional synapse-regulating circRNAs and their predicted miRNA associations.

### circRERE function in (patho-)physiology

Intriguing links between circRERE and neurodegenerative disorders have already been provided, in particular regarding Huntington’s Disease (HD). For example, circRNA microarray profiling performed in a rat PC12-Q74 cell line identified 23 circRNAs as differentially expressed. Strikingly, 16 of the 19 downregulated circRNAs represent circRERE isoforms (Marfil-Marin *et al*., 2021). HD is also characterised by alterations in dopaminergic neurotransmission, which is perturbed in miR-128-3p deficient mice (Tan *et al*., 2013), thereby providing a link between circRERE and miR-128-3p. miR-128-3p further targets HD-associated genes, such as SP1, Huntington Interacting Protein 1 (Hip1) and Htt itself (Kocerha et al., 2014). HD patients develop seizures, pertinent as reduced miR-128-3p is associated with increased neuronal activity (Lanza et al., 2023). Moreover, miR-128-3p levels have also been shown as significantly downregulated in HD patients (Marti et al., 2010), transgenic HD monkeys (Kocerha *et al*., 2014), and transgenic HD mouse models (Lee et al., 2011). Together, these results warrant further investigations into the functional relevance of the circRERE-miR-128-3p interaction for HD development.

Mutations within the RERE gene itself are associated with neurodevelopmental disorders. The “RERE-related neurodevelopmental syndrome” is characterized by global developmental delay, intellectual disability, hypotonia, seizures, and autism spectrum disorder (Niehaus et al., 2022). Accordingly, RERE has been listed as an autism risk gene in the SFARI database (Abrahams et al., 2013) (https://gene.sfari.org/). circRERE levels have also been seen to increase in the hippocampus of a BTBR model of Autism (Gasparini et al., 2020). To what extent circRERE dysregulation, e.g., caused by mutations in non-coding regions of the RERE gene, contribute to autism etiology is an interesting topic for the future. More generally, given the striking overrepresentation of synaptic circRNA host genes in the SFARI database, circRNA dysregulation should be considered as potential pathophysiological mechanism in autism and other synaptopathies.

## Methods

### Primary Cell culture

Primary cortical and hippocampal neuronal cultures were prepared from embryonic day 18 (E18) male and female Sprague-Dawley rats (Janvier Laboratories) as described previously (Schratt et al., 2006). Euthanasia of pregnant rats for removal of embryonic brains was approved by the Veterinary Office of the Canton Zurich, Switzerland, under license ZH027/2021. Dissociated cortical neurons were seeded on poly-L-ornithine plated six-well plates (used for nucleofections), whereas hippocampal neurons were seeded on poly-L-lysine/laminin-coated coverslips in 24-well plates. Except for electrophysiology, all neuron cultures were maintained in Neurobasal-plus (Thermo Fisher Scientific, A3582901) media supplemented with, 2 mM GlutaMAX, 2% B27 100 μg/ml streptomycin, and 100 U/ml penicillin (Invitrogen, Gibco) in an incubator with 5% CO2 at 37°C. Primary hippocampal neurons used for electrophysiology were maintained in Neurobasal-A (Thermo Fisher Scientific, 10888022) media supplemented with 2% B27, 2 mM GlutaMAX, 100 μg/ml streptomycin, and 100 U/ml penicillin (Invitrogen, Gibco).

### Transfection/Nucleofection

All transfections of hippocampal cells were performed using Lipofectamine 2000 (Invitrogen), in duplicate/triplicate wells on DIV8/9 in Neurobasal plus medium (Thermo Fisher Scientific, A3582901), with the exception of electrophysiology experiments. 1 μg of total DNA was transfected per well in a 24-well plate, where empty pcDNA3.1 vector was used to make up the total amount of DNA. Neurons were transfected in Neurobasal plus media in the absence of streptomycin and penicillin for 2 h, replaced with neuron culture media containing ApV (1:1,000) for 45 min, which was washed away and replaced with conditioned media.

Hippocampal neuron transfections for electrophysiology were performed using Lipofectamine 2000 at DIV9 in Neurobasal-A medium (Thermo Fisher, 10888022). 1 μg of total DNA was transfected per well in a 24-well plate, where an empty pcDNA3.1 vector was used to make up the total amount of DNA. Before addition of the lipofectamine/DNA mix, cells were equilibrated in warm Neurobasal-A containing ApV (1:1,000) without penicillin and streptomycin for 30 min in 37°C. Transfection incubation time was reduced to 1.5 h. Cells were then incubated with Neurobasal-A supplemented with ApV for 45 minutes, replaced with conditioned media, and maintained until the day of recording (DIV15/16). Transfections of primary hippocampal neurons utilised 7.5ng of pSUPER constructs, 7.5 pmol/107.5 ng of siRNA, 150ng GFP, 300ng of circRNA/miRNA overexpression constructs. In the case of circRERE Sh, as described in the text, 2.5ng of circRERE1Sh, circRERE2Sh2 and circRERE3Sh pSUPER constructs was utilised instead of 7.5ng of a single shRNA expressing pSUPER plasmid.

Nucleofections were done on cortical neurons using the P3 Primary Cell 4D-Nucleofector X Kit (Lonza, LZ-V4XP-3024), on the day of preparation and dissociation (DIV 0). 4 million dissociated cortical cells were electroporated with 3 μg total DNA per condition with the DC-104 program, seeded in six-well plates in DMEM/ GlutaMAX supplemented with 5% FBS and incubated for 4 h and then replaced with neuron culture media and incubated at 37°C until harvesting. The following amounts of DNA were used for the relevant nucleofections: 2ug pSUPER plasmid or circRNA OE plasmid, 1ug GFP plasmid.

### circRNA reconstruction

Unique circRNA BSJs were reconstructed from RNA-sequencing data from (Colameo *et al*., 2021) using the Deep computational Circular (DCC) RNA Analytics approach/circTest. Available at: https://github.com/dieterich-lab/DCC and https://github.com/dieterich-lab/CircTest.

### RNA Interference

Silencer Select siRNA Custom RNAi Screen (Thermo Fisher) pools of 3x siRNAs against each investigated circRNA BSJ were generated, each staggered by 2-3nt. Sense and antisense sequences for each siRNA provided in supplementary primer table. https://www.thermofisher.com/order/custom-genomic-products/tools/sirna/. pSUPER shRNA-expressing plasmids (pSUPER Basic, Oligoengine, VEC-PBS0001/0002) producing19nt mature shRNA sequences were generated by oligo cloning between BglII and HindIII restriction enzyme sites and verified by sanger sequencing. Sense sequences provided in supplementary primer table.

### circRNA Overexpression

Circular RNA overexpression utilised the Wilusz lab’s pcDNA3.1(+) ZKSCAN1 MCS Exon Vector (Plasmid #69901, Addgene), with the sense sequence of the RERE exons 5-12 (ENSRNOT00000024443.4) cloned between EcoRV and SacII restriction enzyme sites by PCR extension addition of RE sites from randomly primed hippocampal rat cDNA, with the mutant synthesised by Geneart synthesis (Thermo-Fisher). Cortical primary neurons at DIV0 were nucleofected with 2ug of the OE or Mut plasmid and 1ug GFP to confirm transfection efficiency. In primary hippocampal neurons, for synapse density experiments (Figure 7), 300ng OE or Mut plasmid was utilised for lipofection alongside 7.5ng pSUPER shRNA plasmid, 150ng GFP and made to 1ug with empty pcDNA3.1 vector.

### miR-128-2 Overexpression

dsRed-miR-128-2 overexpression construct producing the chimeric intron of miR-128-2 and the control overexpression construct overexpression and control intron (dsRed-β-Actin) were kindly provided by Gregory Wulczyn (Charite Berlin), details provided in (Rehfeld *et al*., 2018).

### Immunocytochemistry, spine density, and image analysis

Stainings were performed on neurons fixed with 4% paraformaldehyde/4% sucrose/PBS. Fixation time was limited to 10-15 min to assure intact post-synaptic compartments. For all imaging experiments, hippocampal cells were transfected on DIV8/9 in duplicate or triplicate wells in 24 well plates. For GluA1 surface staining, live cells were treated with primary antibody at 37°C for 1 h. After washing the cells 4x times with fresh cell media, cells were fixed for 13 min with 4% paraformaldehyde/4% sucrose/PBS and washed with PBS. Coverslips were then transferred to a humidified chamber at room temperature, incubated in secondary antibody in GDB for 1 h, washed with PBS, rinsed briefly with MilliQ water, and mounted onto glass slides for imaging. For PSD-95/Synapsin1 co-staining, following 13-minute 4% paraformaldehyde/4% sucrose/PBS fixation, the coverslips were incubated in GDB for 20 minutes. They were then immediately incubated at room temperature with primary antibodies diluted in GDB in the dark for 2 hours and washed 4x times for 5-10 minutes. Secondary antibodies in GDB were applied for 1.5 hr alongside Hoechst (1:2000). The following primary antibodies were used: Rabbit polyclonal anti-GluA1 (PC246 Calbiochem EMD Biosciences, at final concentrations 2 μg/ml for surface staining), rabbit anti-Synapsin1 (AB1543, 1:1,000; Merck Millipore), and mouse anti-PSD-95 (810401, 1:200; BioLegend). Alexa 488-546– and 647–conjugated secondary antibodies (1:2,000 dilution) were used for detection. Hoechst 33342 (Thermo Fisher) 1:2000 (DNA/Nuclear marker). Images are acquired with a confocal laser scanning microscope (Zeiss, CLSM 880) at 63x unless otherwise stated. Images were processed by Airyscan processing at 6.0 strength 3D, and maximum intensity projections of the Z-stacks were used for signal quantification.PSD-95/Synapsin co-cluster number and surface GluA1 particle number within cell area were analyzed with a custom-made Python script developed by D Colameo and can be added as a Plugin on Fiji (https://github.com/dcolam/Cluster-Analysis-Plugin). Fiji image analysis software available at https://imagej.net/software/fiji/downloads.Transfected GFP plasmid produced a neuronal mask used to determine the inclusion of PSD95/Syn1 Co-Clusters or Glua1 surface puncta within the neuron of interest. The number of puncta/Co-Clusters was then normalized to the area of the GFP mask referred to as “Synapse density” or “GluA1 Surface Puncta per square micron”. Dendritic spines were also detected with our custom Fiji plugin based upon the structural characteristics of spines based off the GFP signal and intensity and the protrusion of the spine relative to its dendrite. Full details available as CFG files for each experiment. In the case of the primary siRNA screen (Figure 2), each biological replicate (N) was spread over 4 different 24-well plates due to the size of the screen. To account for this, each condition was normalized to a control condition on each plate (GFP) and then compared to siControl conditions (also GFP normalized). Immunocytochemistry analysis was performed blinded, primary screen blinding scheme and colour coding noted in extended R markdown.

### circRNA/mRNA FISH and miRNA FISH

Single-molecule (sm) FISH was performed using the ViewRNA Cell Plus kit according to manufacturers protocol (Thermo Fisher - 88-19000-99) with probes for RERE mRNA (vc4-3148005-VCP) and a custom probe against RERE exons 5-10. circRNA FISH for circStau2, circRMST and circHomer1 instead utilised the ViewRNA Cell kit, with signal development with Fast Red Substrate with probe sequences provided in supplementary primer table. For miR-128-3p miRNA FISH, an EDC cross-labelling step was added before the protocol (Thermo Fisher, 22980), using solutions of the Affymetrix QuantiGene ViewRNA miRNA ISH Cell Assay kit and following the manufacturers described steps. An additional Protease treatment was utilised for miRNA FISH; to preserve dendrite morphology, Protease QS was used at a dilution of 1: 10,000 for 45 sec. Rat Camk2a (VC4-15081) acted as a marker for excitatory neurons and allowed for the determination of cellular inclusion of other probe signals as CamkIIa fills the soma compartment and the proximal dendrites. hsa-miR-128-3p viewRNA Cell Plus probe (Assay ID:VM1-10249-VCP) allowed for detection of miR-128-3p signal. Note: miR-128-3p mature sequence is conserved between human, mouse and rat.

### Deprecated Sholl analysis

Images were taken on an Axio Observer Inverted Brightfield Microscope (Zeiss) using a 20x objective. The coverslip was subjected to tiling to reconstruct a high-resolution image of the full coverslip; this allowed for a more rigorous selection of representative cells for the overall coverslip. 200-250 tiles covered the majority of the coverslip, with an image resolution of ∼35000 X ∼23300 pixels, corresponding to an image size of ∼7600 x ∼5000 μm before cell selection. The level of dendritic arborisation of pyramidal neurons was determined by Sholl analysis. This involved ten concentric circles being superimposed around the midpoint of the soma at 20um increments. The number of intersections across each circle was counted, allowing the Sholl profile of the dendritic arbor to be obtained. This process was automated with the FIJI “Deprecated Sholl” programme. Ten neurons for each condition were analysed per independent biological replicate (n=3).

### Adult rat brain tissue collection

Adult Rat (female) was sacrificed by cervical dislocation and various brain tissues collected on an ice-cold glass plate and subsequentially snap-frozen for RNA extraction (Trizol protocol) and gene expression analysis (qPCR) after Superscript III reverse transcription by random priming.

### RNA extraction and quantitative real-time PCR

RNA was isolated using RNA-Solv reagent (Omega Bio-tek) (tissue) or mirVana miRNA extraction kit (Cell culture). Genomic DNA was removed with TURBO DNAse enzyme (Thermo Fisher Scientific). Reverse transcription was performed using either the Taqman MicroRNA Reverse Transcription Kit (Thermo Fisher Scientific) for miRNA detection or the Superscript III reverse transcriptase for mRNA/circRNA detection (Thermo Fisher) with random hexamer priming. qPCR was performed using either Taqman Universal PCR Master Mix (Thermo Fisher Scientific) for microRNA detection or the iTaq SYBR Green Supermix with ROX (Bio-Rad) for mRNA/circRNA, and plates were read on the CFX384 Real-Time System (Bio-Rad). Data were analyzed via the ΔΔCt method and normalized to either U6 (for miRNAs) or GAPDH/Ywhaz/U6 (for mRNAs/circRNAs). mRNA/circRNA convergent (linear mRNA) and divergent (circRNA) primer information is indicated in supplemental primer table. Taqman primers used were (Thermo Fisher Scientific): U6 snRNA (Assay ID: 001973), hsa-miR-128a-3p (Assay ID:002216).

### Luciferase assay miR-128-3p 2x perfect binding site reporters

Luciferase assays were performed using a dual luciferase reporter assay (pmirGLO vector, Promega). 2x perfect binding sites for miR-128-3p were inserted into the firefly 3’UTR between the NheI/SalI restriction enzyme sites. Full sequence in supplementary primer table. Triplicate conditions were transfected in primary HC neurons at DIV9, 50ng of the pMirGlo plasmid of interest alongside 7.5ng pSUPER construct or 300ng miR128 OE construct, 150ng GFP and made up to 1ug with pcDNA3.1. At DIV12/13 cells were lysd in Passive Lysis Buffer (diluted to 1×; Promega) for 15 min, and dual-luciferase assay performed using homemade reagents (as described in (Inouye *et al*., 2022)) on the GloMax Discover GM3000 (Promega).

### RNA Sequencing culturing conditions/RNA Extraction

Cortical neuronal cultures after electroporation at DIV0 were maintained in Neurobasal-plus medium supplemented with 2% B27, 3mM GlutaMAX, 100ug/ml streptomycin and 100u/ml penicillin (Thermo) in a 37°C incubator with 5% CO2 until lysis at DIV5. mirVana miRNA isolation kit with Total RNA Isolation Procedure was utilised for the extraction of total RNA. DNase treatment (Turbo DNase, Thermo Fisher) was performed to remove DNA contamination. The same RNA material was then used for Poly-A and Small RNA sequencing (Novogene) as described.

### mRNA non-directional (polyA) sequencing (Novogene)

RNA sample was used for library preparation using NEB Next® Ultra RNA Library Prep Kit for Illumina®. Indices were included to multiplex multiple samples. Briefly, mRNA was purified from total RNA using poly-T oligo-attached magnetic beads.

After fragmentation, the first strand cDNA was synthesized using random hexamer primers followed by the second strand cDNA synthesis. The library was ready after end repair, A-tailing, adapter ligation, and size selection. After amplification and purification, insert size of the library was validated on an Agilent 2100 and quantified using quantitative PCR (Q-PCR). Libraries were then sequenced on Illumina NovaSeq 6000 with PE150 according to results from library quality control and expected data volume. Quantification performed using salmon 1.8.0 with -- validateMappings on the mRatBN7.2 transcriptome. Surrogate variable analysis was performed with the sva package and two SVs used in the differential expression model. Features were filtered with edgeR::filterByExpr with min.count=20, and differential expression was performed with edgeR’s likelihood ratio test, comparing the knockdown to the two kinds of control samples taken together (the decision to pool the two control groups together was taken after observing the general lack of difference between them). (Patro et al., 2017), (Robinson et al., 2010), (Leek et al., 2012).

### Small RNA Seq (Novogene)

The quality control procedures for total RNA include agarose gel electrophoresis (1%, 180V, 16 min) and Nanodrop Spectrophotometer measurement, which are applied for preliminary monitoring of RNA degradation, concentration, and purity. Agilent 2100 Bioanalyzer is used to test RNA integrity accurate quantification of RNA. After quality control checks, small RNA library was prepared with the NEB Next® Multiplex Small RNA Library Prep Set for Illumina® according to the manufacturer’s instructions.

Briefly, 3’ and 5’ adaptors were ligated to 3’ and 5’ end of small RNA, respectively. Then the first strand cDNA was synthesized after hybridization with reverse transcription primer. The double-stranded cDNA library was generated through PCR enrichment. After purification and size selection, libraries with insertions between 18∼40 bp were ready for sequencing with SE50. The constructed libraries underwent quality controls including assessment of size distribution (Agilent 2100 Bioanalyzer) and molarity (quantified by qPCR). Qualified libraries were sequenced on an Illumina NovaSeq 6000 platform using S4 flow cells (Illumina, USA) using a SE50 bp mode. GEO accession ID: GSE261610. Short RNA analysis was performed with sports 1.0 with -M 1, using the authors’ Rnor6 annotation. Reads were aggregated between genome- and library-aligned, and only miRNAs were considered for downstream analysis. Surrogate variable analysis was performed with the sva package and two SVs used in the differential expression model. Features were filtered with edgeR::filterByExpr with min.count=30, and differential expression was performed as for the poly-A RNA. (Shi et al., 2018)

### GO Term Analysis/Gene Set Enrichment Analysis

Gene Ontology enrichment analysis (GO term) was performed using the TopGo algorithm (v.2.52.0) (Alexa et al., 2006), as described previously in (Soutschek and Schratt, 2023). Available at https://bioconductor.org/packages/release/bioc/html/topGO.html.

Gene set enrichment analysis (GSEA) was conducted with the adaptive multilevel approach from the fgsea package v1.160 (Sergushichev, 2016) against mouse Hallmark and GO terms from the msigdbr package v7.5.1 (Dolgalev, 2022) using the logFC-signed -log10(p-value) of genes passing DEA-filtering (see above) as signal, using the logFC-signed -log10(p-value) of genes passing DEA-filtering (see above) as signal.

### Rolling Circle Amplification

Performed essentially as described in Adult rat cortical RNA was subjected to reverse transcription with random hexamers and PCR-amplified using circRERE2 BSJ-flanking or BSJ-spanning divergent qPCR primers, (Noted in Supplementary Primer list). Running PCR samples on TAE 1.5% Agarose gel electrophoresis revealed the expected size for the respective circRNA species.

### RNase R Treatment**/**qPCR

Total RNA extract from adult rat cortex (2ug) was incubated with 3U/ug of RNase R (or mock-treated) at 37°C for 10 minutes. Subsequently, the RNA was transferred back to ice and a 10% (200ng) spike-in of E. coli total RNA was added. The RNA was re-extracted with acidic phenol-chloroform and ethanol-precipitated. The RNA concentration of the mock-treated sample was determined, and 1 µg was used for reverse transcription with Superscript III (Invitrogen) as described above, the same volume of the RNase R-treated RNA was used for reverse transcription. The E. coli spike-in was used for normalization in the qPCR with CysG primers.

### Electrophysiology

Whole-cell patch-clamp recordings were performed on an upright microscope (Olympus BX51WI) at room temperature. Data were collected with an Axon MultiClamp 700B amplifier and a Digidata 1550B digitizer and analysed with pClamp11 software (all from Molecular Devices). Recording pipettes were pulled from borosilicate capillary glass (GC150F-10; Harvard Apparatus) with a DMZ-Universal-Electrode-Puller (Zeitz) and had resistances between 3 and 4 MW.

Miniature EPSCs (mEPSCs) were recorded from primary cultured hippocampal neurons on DIV15-16 after transfection in Neurobasal-A medium (10888022; Thermo Fisher Scientific) on DIV9. The extracellular solution (ACSF) was composed of (in mM) 140 NaCl, 2.5 KCl, 10 Hepes, 2 CaCl2, 2 MgCl2, 10 glucose (adjusted to pH 7.3 with NaOH), the intracellular solution of (in mM) 125 K-gluconate, 20 KCL, 0.5 EGTA, 10 Hepes, 4 Mg-ATP, 0.3 GTP, and 10 Na2-phosphocreatine (adjusted to pH 7.3 with KOH). For mEPSCs, 1 μM TTX and 1 μM Gabazine were added to the extracellular solution to block action-potential driven glutamate release and GABAergic synaptic transmission, respectively. Cells were held at −60 mV. The sampling frequency was 5 kHz, and the filter frequency 2 kHz. Series resistance was monitored, and recordings were discarded if the series resistance changed significantly (≥10%) or exceeded 22 MΩ.

### ScanmiR

miRNA binding site prediction on reconstructed circRNA isoforms utilised the ScaMiR webtool version 1.5.2 https://ethz-ins.org/scanMiR/ based upon the Rat refseq Rn7 miRNA collection using default parameters as described (Soutschek *et al*., 2022b).

### EnrichmiR

enrichMiR analyses and cumulative distribution (CD) plots were generated using the enrichMiR package. A CD-plot was generated on the scanMiR annotation using the option to split by site affinity. Enrichment Plot utilises aggregated statistical model with a significance-weighted average of the scaled enrichment scores. Repression scores were inverted, scores were scaled by unit variance (across miRNAs), for each miRNA the tests were weighted by the significance quantile within the test and then averaged across tests. Further information available at enrichMiR publication. EnrichMir version 0.99.28 https://ethz-ins.org/enrichMiR/ (Soutschek *et al*., 2022a).

### Statistical analyses

For the primary circRNA siRNA screen (Fig. 2), outputs were first normalized to the GFP condition, that served as a transfection control, and then log-transformed to decrease variance. Linear mixed effects models (using lme4 package) for nested data were used to model the relationship between siRNA pools and synapse co-cluster, spine density and volume. Post hoc multiple comparisons against a single siControl1, were conducted with the emmeans package using Dunnett’s procedure. The same model specifications were used for synapse density, spine density and volume analyses (example of model specification provided below). A similar approach was used to analyse synapse density upon shRNA mediated knockdown of circRERE isoforms (Fig. 3) and GluA1 puncta density (Fig. 4).

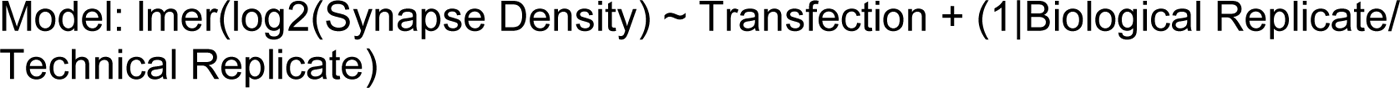

For the analysis of synapse density with circREREsh and miR-128-3p overexpression (Fig. 6), a linear mixed effects model with pairwise post hoc comparisons and multiple-testing correction was used to calculate p-values, where the biological replicates are defined as a random effect.

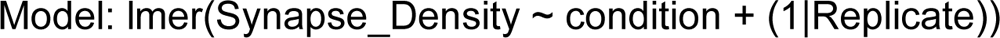

To analyse synapse density upon transfection of combinatorial shRNA and miRNA overexpression constructs (Fig. 7), additional predictors were added to the general linear model to correct for the variance generated by mosaic plasmid expression and overexpression in an interaction test. Model specified below; for complete model matrix, please reference provided scripts.

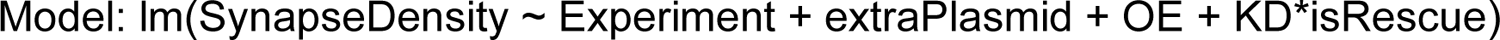

Plots were generated in R using various packages, including ggplot2 detailed in R markdowns. ImageJ/FIJI was used to adjust contrasts in microscope images as well as to insert scale bars.

Graphpad Prism 9 was utilised for the plotting and statistical analysis of electrophysiology recordings, supplemental luciferase assays and qPCRs utilising either 1-way ANOVA or unpaired T-Tests.

## Author Contributions

DK designed project, prepared RNA samples, microscopy experiments, cell culture, treatments, stainings, RT-qPCR experiments, prepared figures and wrote the original manuscript draft. SB performed optimization of circRNA RNAi, circFISH and primary knockdown screen. JW performed patch-clamp recording of primary HC cultures. PN performed statistical analysis and generated plots for all histology experiments. PLG performed RNA-seq analysis, statistical analysis of the total circRNA population in compartmentalized neurons and overlap with published datasets. CD reconstructed circRNA BSJs from RNA-seq data and performed statistical assessment. GS supervised the project, coordinated the collaboration, and wrote the manuscript.

## Acknowledgments

We greatly acknowledge the excellent technical assistance provided by Tatjana Wüst, Cristina Furler and Roberto Fiore. We further thank David Colameo for the establishment and maintenance of FIJI-based image analysis pipelines. We would like to thank visiting students: Alexandra Huber and Martin Breu for assistance with extended morphological analyses, Yu Wang for assistance with miRNA FISH analysis, and Connor Bitter and Julianna Strother for testing of miR-128OE constructs. We thank Gregory Wulczyn (Charite Berlin) for the gift of Mir-128-2-OE constructs. D.K. was supported by a PhD fellowship from the SNSF, NCCR RNA & Disease. G.S. received funding from the Deutsche Forschungsgemeinschaft (DFG; grant SPP1738 (SCHR1136/4-2)) and the Swiss National Science Foundation (SNSF; grants NeuroCirc ([IZSTZ0_216044](https://www.mysnf.ch/login.aspx?returnurl=%2fgrants%2fgrant.aspx%3fid%3d16a5be54-3c5d-472e-8a53-69c5359985dc)) and ALTRUISM ([32NE30_189486](https://www.mysnf.ch/login.aspx?returnurl=%2fgrants%2fgrant.aspx%3fid%3d842a6ab6-7220-4ac6-b53d-5f816c4bf341))). C.D. received funding from the Deutsche Forschungsgemeinschaft (DFG; grant SPP1738 (DI 1501/5-2).

## Data availability

RNA sequencing data has been deposited to Gene Expression Omnibus (GEO, Poly-A sequencing Fig. 5A, Small-RNA sequencing Fig 6A, accession number GSE261610).

## References

Abrahams, B., Arking, D., Campbell, D., Mefford, H., Morrow, E., Weiss, L., Menashe, I., Wadkins, T., Banerjee-Basu, S., and Packer, A. (2013). SFARI Gene 2.0: a community-driven knowledgebase for the autism spectrum disorders (ASDs). Molecular Autism 4.

Alexa, A., Rahnenfuhrer, J., and Lengauer, T. (2006). Improved scoring of functional groups from gene expression data by decorrelating GO graph structure. Bioinformatics 22, 1600–1607. 10.1093/bioinformatics/btl140.

Ashwal-Fluss, R., Meyer, M., Pamudurti, N.R., Ivanov, A., Bartok, O., Hanan, M., Evantal, N., Memczak, S., Rajewsky, N., and Kadener, S. (2014). CircRNA Biogenesis competes with Pre-mRNA splicing. Molecular Cell. Elsevier.

Bicker, S., Khudayberdiev, S., Weiss, K., Zocher, K., Baumeister, S., and Schratt, G. (2013). The DEAH-box helicase DHX36 mediates dendritic localization of the neuronal precursor-microRNA-134. Genes Dev 27, 991–996. 10.1101/gad.211243.112.

Chi, S.W., Zang, J.B., Mele, A., and Darnell, R.B. (2009). Argonaute HITS-CLIP decodes microRNA-mRNA interaction maps. Nature 460, 479–486. 10.1038/nature08170.

Colameo, D., Rajman, M., Soutschek, M., Bicker, S., von Ziegler, L., Bohacek, J., Winterer, J., Germain, P.L., Dieterich, C., and Schratt, G. (2021). Pervasive compartment-specific regulation of gene expression during homeostatic synaptic scaling. EMBO Rep 22, e52094. 10.15252/embr.202052094.

Das, A., Rout, P.K., Gorospe, M., and Panda, A.C. (2019). Rolling Circle cDNA Synthesis Uncovers Circular RNA Splice Variants. International Journal of Molecular Sciences.

Daswani, R., Gilardi, C., Soutschek, M., Nanda, P., Weiss, K., Bicker, S., Fiore, R., Dieterich, C., Germain, P.L., Winterer, J., and Schratt, G. (2022). MicroRNA-138 controls hippocampal interneuron function and short-term memory in mice. Elife 11. 10.7554/eLife.74056.

de la Mata, M., Gaidatzis, D., Vitanescu, M., Stadler, M.B., Wentzel, C., Scheiffele, P., Filipowicz, W., and Grosshans, H. (2015). Potent degradation of neuronal miRNAs induced by highly complementary targets. EMBO Rep 16, 500–511. 10.15252/embr.201540078.

Dell’Orco, M., Oliver, R.J., and Perrone-Bizzozero, N. (2020). HuD Binds to and Regulates Circular RNAs Derived From Neuronal Development- and Synaptic Plasticity-Associated Genes. Frontiers in Genetics.

Diering, G.H., and Huganir, R.L. (2018). The AMPA Receptor Code of Synaptic Plasticity. Neuron 100, 314–329. 10.1016/j.neuron.2018.10.018.

Dolgalev, I. (2022). msigdbr: MSigDB gene sets for multiple organisms in a tidy data format. R package version 7.5. 1.9001.

Dong, X., Bai, Y., Liao, Z., Gritsch, D., Liu, X., Wang, T., Borges-Monroy, R., Ehrlich, A., Serrano, G.E., Feany, M.B., et al. (2023). Circular RNAs in the human brain are tailored to neuron identity and neuropsychiatric disease. Nat Commun 14, 5327. 10.1038/s41467-023-40348-0.

Du, M., Wu, C., Yu, R., Cheng, Y., Tang, Z., Wu, B., Fu, J., Tan, W., Zhou, Q., Zhu, Z., et al. (2022). A novel circular RNA, circIgfbp2, links neural plasticity and anxiety through targeting mitochondrial dysfunction and oxidative stress-induced synapse dysfunction after traumatic brain injury. Mol Psychiatry 27, 4575–4589. 10.1038/s41380-022-01711-7.

Dube, U., Del-Aguila, J.L., Li, Z., Budde, J.P., Jiang, S., Hsu, S., Ibanez, L., Fernandez, M.V., Farias, F., Norton, J., et al. (2019). An atlas of cortical circular RNA expression in Alzheimer disease brains demonstrates clinical and pathological associations. Nature Neuroscience.

Epple, R., Kruger, D., Berulava, T., Brehm, G., Ninov, M., Islam, R., Koster, S., and Fischer, A. (2021). The Coding and Small Non-coding Hippocampal Synaptic RNAome. Mol Neurobiol 58, 2940–2953. 10.1007/s12035-021-02296-y.

Fernandez, E., Li, K.W., Rajan, N., De Rubeis, S., Fiers, M., Smit, A.B., Achsel, T., and Bagni, C. (2015). FXR2P Exerts a Positive Translational Control and Is Required for the Activity-Dependent Increase of PSD95 Expression. J Neurosci 35, 9402–9408. 10.1523/JNEUROSCI.4800-14.2015.

Franzoni, E., Booker, S.A., Parthasarathy, S., Rehfeld, F., Grosser, S., Srivatsa, S., Fuchs, H.R., Tarabykin, V., Vida, I., and Wulczyn, F.G. (2015). miR-128 regulates neuronal migration, outgrowth and intrinsic excitability via the intellectual disability gene Phf6. Elife 4. 10.7554/eLife.04263.

Fuchs Wightman, F., Lukin, J., Giusti, S.A., Soutschek, M., Bragado, L., Pozzi, B., Pierelli, M.L., Gonzalez, P., Fededa, J.P., Schratt, G., et al. (2024). Influence of RNA circularity on Target RNA-Directed MicroRNA Degradation. Nucleic Acids Res. 10.1093/nar/gkae094.

Gardoni, F., Saraceno, C., Malinverno, M., Marcello, E., Verpelli, C., Sala, C., and Di Luca, M. (2012). The neuropeptide PACAP38 induces dendritic spine remodeling through ADAM10-N-cadherin signaling pathway. J Cell Sci 125, 1401–1406. 10.1242/jcs.097576.

Gasparini, S., Del Vecchio, G., Gioiosa, S., Flati, T., Castrignano, T., Legnini, I., Licursi, V., Ricceri, L., Scattoni, M.L., Rinaldi, A., et al. (2020). Differential Expression of Hippocampal Circular RNAs in the BTBR Mouse Model for Autism Spectrum Disorder. Mol Neurobiol 57, 2301–2313. 10.1007/s12035-020-01878-6

Gokool, A., Anwar, F., and Voineagu, I. (2019). The Landscape of Circular RNA Expression in the Human Brain. Biological Psychiatry. Elsevier Inc.

Gomes-Duarte, A., Veno, M.T., de Wit, M., Senthilkumar, K., Broekhoven, M.H., van den Herik, J., Heeres, F.R., van Rossum, D., Rybiczka-Tesulov, M., Legnini, I., et al. (2022). Expression of Circ_Satb1 Is Decreased in Mesial Temporal Lobe Epilepsy and Regulates Dendritic Spine Morphology. Front Mol Neurosci 15, 832133. 10.3389/fnmol.2022.832133.

Guo, J., Agarwal, V., Guo, H., and Bartel, D. (2014). Expanded identification and characterization of mammalian circular RNAs. Genome Biology 15 (409).

Hanan, M., Simchovitz, A., Yayon, N., Vaknine, S., Cohen-Fultheim, R., Karmon, M., Madrer, N., Rohrlich, T.M., Maman, M., Bennett, E.R., et al. (2020). A Parkinson’s disease CircRNAs Resource reveals a link between circSLC8A1 and oxidative stress. EMBO Mol Med 12, e11942. 10.15252/emmm.201911942.

Hansen, T.B., Jensen, T.I., Clausen, B.H., Bramsen, J.B., Finsen, B., Damgaard, C.K., and Kjems, J. (2013). Natural RNA circles function as efficient microRNA sponges. Nature. Nature Publishing Group.

Hollensen, A.K., Thomsen, H.S., Lloret-Llinares, M., Kamstrup, A.B., Jensen, J.M., Luckmann, M., Birkmose, N., Palmfeldt, J., Jensen, T.H., Hansen, T.B., and Damgaard, C.K. (2020). circZNF827 nucleates a transcription inhibitory complex to balance neuronal differentiation. Elife 9. 10.7554/eLife.58478.

Inouye, M.O., Colameo, D., Ammann, I., Winterer, J., and Schratt, G. (2022). miR-329- and miR-495-mediated Prr7 down-regulation is required for homeostatic synaptic depression in rat hippocampal neurons. Life Sci Alliance 5. 10.26508/lsa.202201520.

Jara, C.A.C., Kim, S.J., Thomas, G., Farsi, Z., Zolotarov, G., Georgii, E., Woehler, A., Piwecka, M., and Rajewsky, N. (2023). miR-7 controls glutamatergic transmission and neuronal connectivity in a Cdr1as dependent manner. bioRxiv. 10.1101/2023.01.26.525729.

Jeck, W.R., Sorrentino, J.A., Wang, K., Slevin, M.K., Burd, C.E., Liu, J., Marzluff, W.F., and Sharpless, N.E. (2013). Circular RNAs are abundant, conserved, and associated with ALU repeats. RNA 19, 141–157. 10.1261/rna.035667.112.

Jonas, P. (2000). The Time Course of Signaling at Central Glutamatergic Synapses. News Physiol. Sci. 15.

Kleaveland, B., Shi, C.Y., Stefano, J., and Bartel, D.P. (2018). A Network of Noncoding Regulatory RNAs Acts in the Mammalian Brain. Cell 174, 350–362 e317. 10.1016/j.cell.2018.05.022.

Kocerha, J., Xu, Y., Prucha, M., Zhao, D., and Chan, A. (2014). microRNA-128a dysregulation in transgenic Huntington’s disease monkeys. Molecular Brain 7:46.

Kramer, M.C., Liang, D., Tatomer, D.C., Gold, B., March, Z.M., Cherry, S., and Wilusz, J.E. (2015). Combinatorial control of Drosophila circular RNA expression by intronic repeats, hnRNPs, and SR proteins. Genes and Development.

Kristensen, L.S., Andersen, M.S., Stagsted, L.V.W., Ebbesen, K.K., Hansen, T.B., and Kjems, J. (2019). The biogenesis, biology and characterization of circular RNAs. Nature Reviews Genetics.

Lanza, M., Cuzzocrea, S., Oddo, S., Esposito, E., and Casili, G. (2023). The Role of miR-128 in Neurodegenerative Diseases. Int J Mol Sci 24. 10.3390/ijms24076024.

Lee, S.T., Chu, K., Im, W.S., Yoon, H.J., Im, J.Y., Park, J.E., Park, K.H., Jung, K.H., Lee, S.K., Kim, M., and Roh, J.K. (2011). Altered microRNA regulation in Huntington’s disease models. Exp Neurol 227, 172–179. 10.1016/j.expneurol.2010.10.012.

Leek, J.T., Johnson, W.E., Parker, H.S., Jaffe, A.E., and Storey, J.D. (2012). The sva package for removing batch effects and other unwanted variation in high-throughput experiments. Bioinformatics 28, 882–883. 10.1093/bioinformatics/bts034.

Li, S., Li, X., Xue, W., Zhang, L., Yang, L.Z., Cao, S.M., Lei, Y.N., Liu, C.X., Guo, S.K., Shan, L., et al. (2021). Screening for functional circular RNAs using the CRISPR-Cas13 system. Nat Methods 18, 51–59. 10.1038/s41592-020-01011-4.

Liu, C.X., and Chen, L.L. (2022). Circular RNAs: Characterization, cellular roles, and applications. Cell 185, 2016–2034. 10.1016/j.cell.2022.04.021.

Liu, L., Neve, M., Perlaza-Jimenez, L., Hawdon, A., Conn, S.J., Zenker, J., Tamayo, P., Goodall, G.J., and Rosenbluh, J. (2022). Systematic loss of function screens identify pathway specific functional circular RNAs. bioRxiv. 10.1101/2022.10.22.513321.

Lu, W., Shi, Y., Jackson, A.C., Bjorgan, K., During, M.J., Sprengel, R., Seeburg, P.H., and Nicoll, R.A. (2009). Subunit composition of synaptic AMPA receptors revealed by a single-cell genetic approach. Neuron 62, 254–268. 10.1016/j.neuron.2009.02.027.

Malinverno, M., Carta, M., Epis, R., Marcello, E., Verpelli, C., Cattabeni, F., Sala, C., Mulle, C., Di Luca, M., and Gardoni, F. (2010). Synaptic localization and activity of ADAM10 regulate excitatory synapses through N-cadherin cleavage. J Neurosci 30, 16343–16355. 10.1523/JNEUROSCI.1984-10.2010.

Marfil-Marin, E., Santamaria-Olmedo, M., PerezGrovas-Saltijeral, A., Valdes-Flores, M., Ochoa-Morales, A., Jara-Prado, A., Sevilla-Montoya, R., Camacho-Molina, A., and Hidalgo-Bravo, A. (2021). circRNA Regulates Dopaminergic Synapse, MAPK, and Long-term Depression Pathways in Huntington Disease. Mol Neurobiol 58, 6222–6231. 10.1007/s12035-021-02536-1.

Marti, E., Pantano, L., Banez-Coronel, M., Llorens, F., Minones-Moyano, E., Porta, S., Sumoy, L., Ferrer, I., and Estivill, X. (2010). A myriad of miRNA variants in control and Huntington’s disease brain regions detected by massively parallel sequencing. Nucleic Acids Res 38, 7219–7235. 10.1093/nar/gkq575.

McSweeney, K.M., Gussow, A.B., Bradrick, S.S., Dugger, S.A., Gelfman, S., Wang, Q., Petrovski, S., Frankel, W.N., Boland, M.J., and Goldstein, D.B. (2016). Inhibition of microRNA 128 promotes excitability of cultured cortical neuronal networks. Genome Res 26, 1411–1416. 10.1101/gr.199828.115.

Memczak, S., Jens, M., Elefsinioti, A., Torti, F., Krueger, J., Rybak, A., Maier, L., Mackowiak, S.D., Gregersen, L.H., Munschauer, M., et al. (2013). Circular RNAs are a large class of animal RNAs with regulatory potency. Nature. Nature Publishing Group.

Ngo, L.H., Bert, A.G., Dredge, B.K., Williams, T., Murphy, V., Li, W., Hamilton, W.B., Carey, K.T., Toubia, J., Pillman, K.A., et al. (2024). Nuclear export of circular RNA. Nature. 10.1038/s41586-024-07060-5.

Niehaus, A.D., Kim, J., and Manning, M.A. (2022). Phenotypic variability in RERE-related disorders and the first report of an inherited variant. Am J Med Genet A 188, 3358–3363. 10.1002/ajmg.a.62952.

Pamudurti, N.R., Bartok, O., Jens, M., Chekulaeva, M., Rajewsky, N., Kadener, S., Pamudurti, N.R., Bartok, O., Jens, M., Ashwal-fluss, R., et al. (2017). Translation of CircRNAs Article Translation of CircRNAs. Molecular Cell. Elsevier Inc.

Paradis, S., Harrar, D.B., Lin, Y., Koon, A.C., Hauser, J.L., Griffith, E.C., Zhu, L., Brass, L.F., Chen, C., and Greenberg, M.E. (2007). An RNAi-based approach identifies molecules required for glutamatergic and GABAergic synapse development. Neuron 53, 217–232. 10.1016/j.neuron.2006.12.012.

Patro, R., Duggal, G., Love, M.I., Irizarry, R.A., and Kingsford, C. (2017). Salmon provides fast and bias-aware quantification of transcript expression. Nat Methods 14, 417–419. 10.1038/nmeth.4197.

Piras, R., Ko, E.Y., Barrett, C., De Simone, M., Lin, X., Broz, M.T., Tessaro, F.H.G., Castillo-Martin, M., Cordon-Cardo, C., Goodridge, H.S., et al. (2022). circCsnk1g3- and circAnkib1-regulated interferon responses in sarcoma promote tumorigenesis by shaping the immune microenvironment. Nat Commun 13, 7243. 10.1038/s41467-022-34872-8.

Piwecka, M., Glažar, P., Hernandez-Miranda, L.R., Memczak, S., Wolf, S.A., Rybak-Wolf, A., Filipchyk, A., Klironomos, F., Jara, C.A.C., Fenske, P., et al. (2017). Loss of a mammalian circular RNA locus causes miRNA deregulation and affects brain function. Science.

Qin, S., Zhang, Q., Xu, Y., Ma, S., Wang, T., Huang, Y., and Ju, S. (2022). m(6)A-modified circRNAs: detections, mechanisms, and prospects in cancers. Mol Med 28, 79. 10.1186/s10020-022-00505-5.

Rehfeld, F., Maticzka, D., Grosser, S., Knauff, P., Eravci, M., Vida, I., Backofen, R., and Wulczyn, F.G. (2018). The RNA-binding protein ARPP21 controls dendritic branching by functionally opposing the miRNA it hosts. Nat Commun 9, 1235. 10.1038/s41467-018-03681-3.

Robinson, M.D., McCarthy, D.J., and Smyth, G.K. (2010). edgeR: a Bioconductor package for differential expression analysis of digital gene expression data. Bioinformatics 26, 139–140. 10.1093/bioinformatics/btp616.

Rybak-Wolf, A., Stottmeister, C., Glažar, P., Jens, M., Pino, N., Hanan, M., Behm, M., Bartok, O., Ashwal-Fluss, R., Herzog, M., et al. (2014). Circular RNAs in the Mammalian Brain Are Highly Abundant, Conserved, and Dynamically Expressed. Molecular Cell.

Schratt, G.M., Tuebing, F., Nigh, E.A., Kane, C.G., Sabatini, M.E., Kiebler, M., and Greenberg, M.E. (2006). A brain-specific microRNA regulates dendritic spine development. Nature.

Seeler, S., Andersen, M.S., Sztanka-Toth, T., Rybiczka-Tesulov, M., van den Munkhof, M.H., Chang, C.C., Maimaitili, M., Veno, M.T., Hansen, T.B., Pasterkamp, R.J., et al. (2023). A Circular RNA Expressed from the FAT3 Locus Regulates Neural Development. Mol Neurobiol 60, 3239–3260. 10.1007/s12035-023-03253-7.

Selcher, J.C., Xu, W., Hanson, J.E., Malenka, R.C., and Madison, D.V. (2012). Glutamate receptor subunit GluA1 is necessary for long-term potentiation and synapse unsilencing, but not long-term depression in mouse hippocampus. Brain Res 1435, 8–14. 10.1016/j.brainres.2011.11.029.

Sergushichev, A.A. (2016). An algorithm for fast preranked gene set enrichment analysis using cumulative statistic calculation. bioRxiv, 060012. 10.1101/060012.

Shi, J., Ko, E.A., Sanders, K.M., Chen, Q., and Zhou, T. (2018). SPORTS1.0: A Tool for Annotating and Profiling Non-coding RNAs Optimized for rRNA- and tRNA-derived Small RNAs. Genomics Proteomics Bioinformatics 16, 144–151. 10.1016/j.gpb.2018.04.004.

Shvarts-Serebro, I., Sheinin, A., Gottfried, I., Adler, L., Schottlender, N., Ashery, U., and Barak, B. (2021). miR-128 as a Regulator of Synaptic Properties in 5xFAD Mice Hippocampal Neurons. J Mol Neurosci 71, 2593–2607. 10.1007/s12031-021-01862-2.

Siegel, G., Obernosterer, G., Fiore, R., Oehmen, M., Christensen, M., Khudayberdiev, S., Leuschner, P.F., Clara, J.L., Kane, C., Hübel, K., et al. (2009). A functional screen implicates microRNA-138-dependent regulation of the depalmitoylation enzyme APT1 in dendritic spine morphogenesis. Nat. Cell Biol.

Soutschek, M., Bianco, A.L., Galkin, S., Wüst, T., Colameo, D., Germade, T., Gross, F., von Ziegler, L., Bohacek, J., Germain, P.-L., et al. (2023). A human-specific microRNA controls the timing of excitatory synaptogenesis. bioRxiv. 10.1101/2023.10.04.560889.

Soutschek, M., Germade, T., Germain, P.L., and Schratt, G. (2022a). enrichMiR predicts functionally relevant microRNAs based on target collections. Nucleic Acids Res 50, W280–W289. 10.1093/nar/gkac395.

Soutschek, M., Gross, F., Schratt, G., and Germain, P.L. (2022b). scanMiR: a biochemically based toolkit for versatile and efficient microRNA target prediction. Bioinformatics 38, 2466–2473. 10.1093/bioinformatics/btac110.

Soutschek, M., and Schratt, G. (2023). Non-coding RNA in the wiring and remodeling of neural circuits. Neuron 111, 2140–2154. 10.1016/j.neuron.2023.04.031.

Tan, C.L., Plotkin, J.L., Veno, M.T., von Schimmelmann, M., Feinberg, P., Mann, S., Handler, A., Kjems, J., Surmeier, D.J., O’Carroll, D., et al. (2013). MicroRNA-128 governs neuronal excitability and motor behavior in mice. Science 342, 1254–1258. 10.1126/science.1244193.

Wu, W., Zhao, F., and Zhang, J. (2024). circAtlas 3.0: a gateway to 3 million curated vertebrate circular RNAs based on a standardized nomenclature scheme. Nucleic Acids Res 52, D52–D60. 10.1093/nar/gkad770.

Xu, C., and Zhang, J. (2021). Mammalian circular RNAs result largely from splicing errors. Cell Rep 36, 109439. 10.1016/j.celrep.2021.109439.

Xu, K., Zhang, Y., Xiong, W., Zhang, Z., Wang, Z., Lv, L., Liu, C., Hu, Z., Zheng, Y.T., Lu, L., et al. (2020). CircGRIA1 shows an age-related increase in male macaque brain and regulates synaptic plasticity and synaptogenesis. Nat Commun 11, 3594. 10.1038/s41467-020-17435-7.

You, X., Hou, J., Sambandan, S., Schuman, E.M., Wang, M., Tushev, G., Babic, A., Will, T., Chen, T., Liu, H., et al. (2015). Neural circular RNAs are derived from synaptic genes and regulated by development and plasticity. Nature Neuroscience.

Zajaczkowski, E.L., Zhao, Q., Liau, W.S., Gong, H., Madugalle, S.U., Periyakaruppiah, A., Leighton, L.J., Musgrove, M., Ren, H., Davies, J., et al. (2023). Localised Cdr1as activity is required for fear extinction memory. Neurobiol Learn Mem 203, 107777. 10.1016/j.nlm.2023.107777.

Zheng, Q., Bao, C., Guo, W., Li, S., Chen, J., Chen, B., Luo, Y., Lyu, D., Li, Y., Shi, G., et al. (2016). Circular RNA profiling reveals an abundant circHIPK3 that regulates cell growth by sponging multiple miRNAs. Nature Communications. Nature Publishing Group.

Zimmerman, A.J., Hafez, A.K., Amoah, S.K., Rodriguez, B.A., Dell’Orco, M., Lozano, E., Hartley, B.J., Alural, B., Lalonde, J., Chander, P., et al. (2020). A psychiatric disease-related circular RNA controls synaptic gene expression and cognition. Molecular Psychiatry.

