## Supplementary figures and methods for "A functional screen uncovers circular RNAs regulating excitatory synaptogenesis in hippocampal neurons"

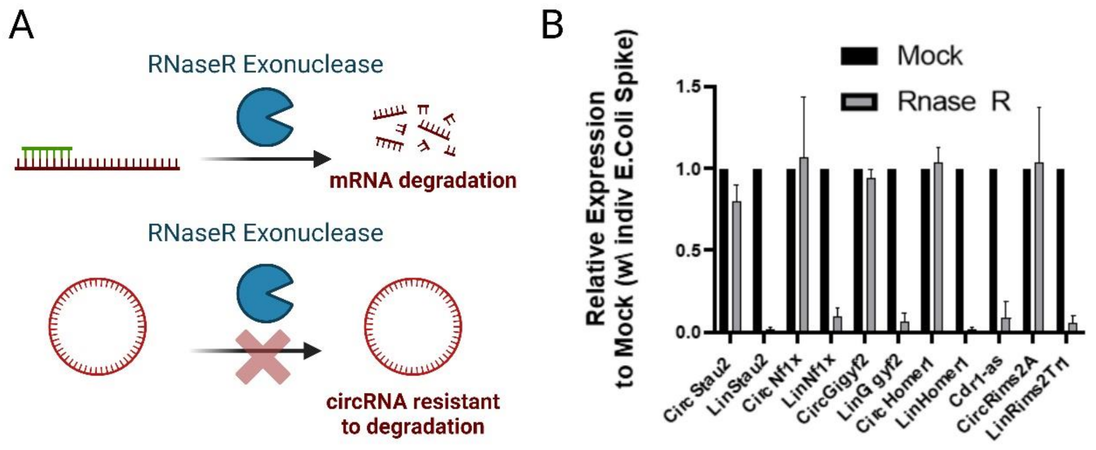


**Suppl. Fig 1: Characterisation of the circRNA population in compartmentalised primary rat hippocampal neurons**

A. Schematic of Rnase R preferential degradation of linear RNA species, while circular RNAs are more RNase R resistant. B. RNase R treatment of purified RNA from adult rat hippocampus and RT-qPCR of selected circRNA candidates and their linear mRNAs. Cdr1-as is only circRNA displaying notable RNase R sensitivity. N=3 biological replicates. CysG normalization from E.Coli spike-in.

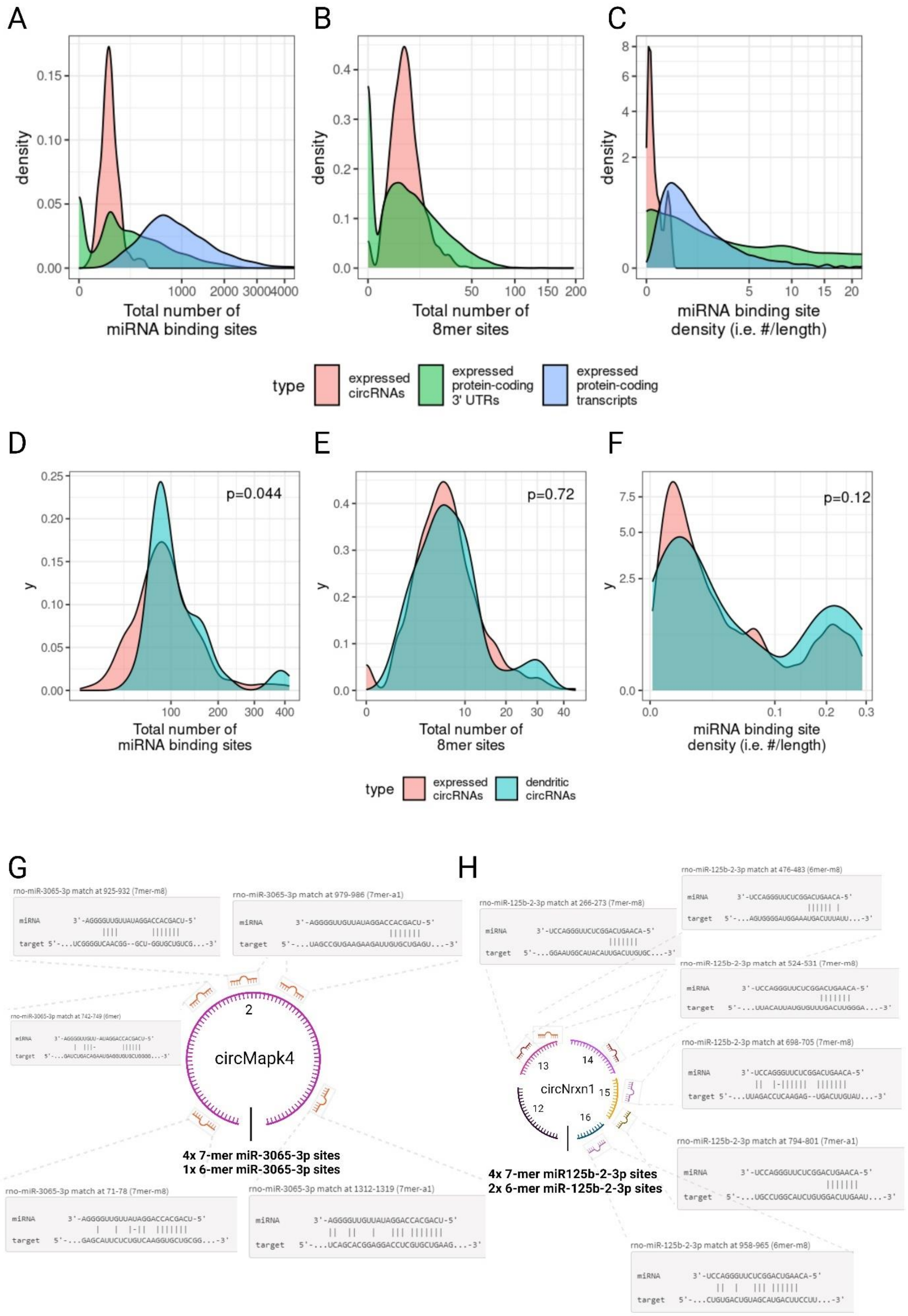
A
B
C
density
Total number of
miRNA binding sites
Total number of
8mer sites
miRNA binding site
density (i.e. #/length)
type
expressed
circRNAs
expressed
protein-coding
3' UTRs
expressed
protein-coding
transcripts
D
E
F
p=0.044
p=0.72
p=0.12
y
type
expressed
circRNAs
dendritic
circRNAs
G
H
circMapk4
4x 7-mer miR-3065-3p sites
1x 6-mer miR-3065-3p sites
circNrxn1
4x 7-mer miR125b-2-3p sites
2x 6-mer miR-125b-2-3p sites

**Suppl. Fig 2: miRNA binding site analysis of circRNA population**

Comparison of the total number and density of indicated miRNA site types in Process-enriched circRNAs (red), 3'UTRs of protein-coding mRNAs (green) or entire protein-coding transcripts (blue). A. Indicating the distribution/density of total miRNA binding sites. B. Indicating the distribution/density of 8mer sites. C. Indicating the relative density of miRNA binding sites normalized to species length. Comparison of the total number and density of indicated miRNA site types in all expressed circRNAs (red) and process-enriched circRNAs (blue). D. Indicating the frequency of total number of miRNA binding sites. E. Indicating the frequency of total number of 8mer sites. F. Indicating the frequency of specific miRNA binding site density. G. ScanMiR miRNA binding site prediction analysis of dendritically enriched circRNA candidate circMapk4 illustrating 5 predicted miR-3065-3p binding sites. H. ScanMiR analysis of circNrxn1, illustrating 6 predicted miR-125b-2-3p binding sites.

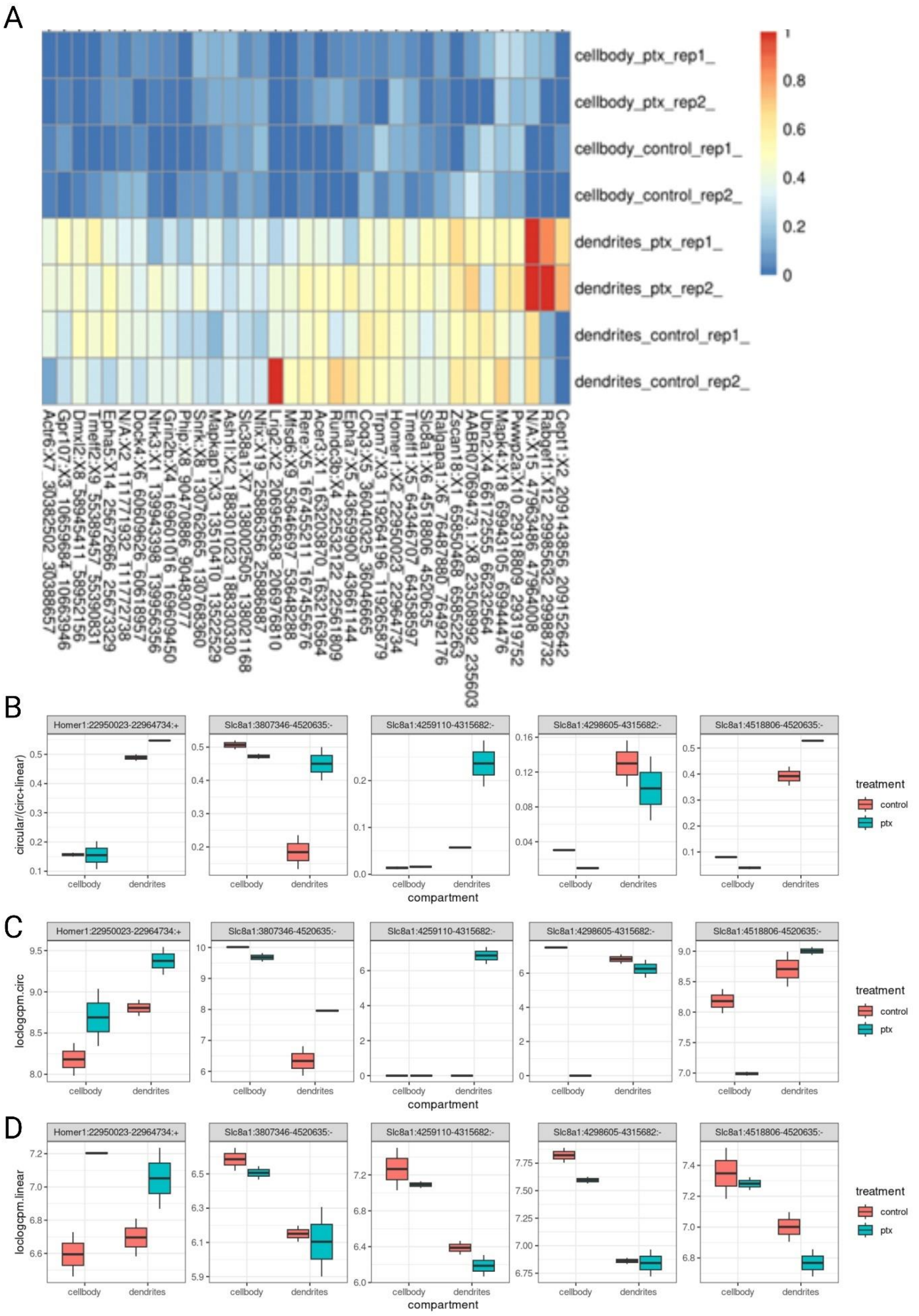
A
cellbody_ptx_rep1_
cellbody_ptx_rep2_
cellbody_control_rep1_
cellbody_control_rep2_
dendrites_ptx_rep1_
dendrites_ptx_rep2_
dendrites_control_rep1_
dendrites_control_rep2_
1
0.8
0.6
0.4
0.2
0
Actr6:X7_30382502_30388657
Gpr107:X3_10659684_10663946
Dmxl2:X8_58945411_58952156
Tmeff2:X9_55389457_55390831
Epha5:X14_25672666_25673329
N/A:X2_111771932_111772738
Dock4:X6_60609626_60618957
Ntrk3:X1_139943398_139956356
Grin2b:X4_169601016_169609450
Phip:X8_90470886_90483077
Snrk:X8_130762665_130768360
Mapkap1:X3_13510410_13522529
Ash1l:X2_188301023_188330330
Slc38a1:X7_138002505_138021168
Nfix:X19_25886356_25886887
Lrig2:X2_206956638_206976810
Mfsd6:X9_53646697_53648288
Rere:X5_167455211_167455676
Acer3:X1_163203870_163216364
Rundc3b:X4_22532122_22561809
Epha7:X5_43659900_43661144
Coq3:X5_36040325_36046665
Trpm7:X3_119264196_119265879
Homer1:X2_22950023_22964734
Tmeff1:X5_64346707_64358597
Slc8a1:X6_4518806_4520635
Ralgapa1:X6_76487880_76492176
Zscan18:X1_65850468_65852263
AABR07069473.1:X8_23508992_23556603
Ubn2:X4_66172555_66232564
Mapk4:X18_69943105_69944476
Pwwp2a:X10_29318809_29319752
N/A:X15_47963486_47964008
Rabgef1:X12_29985632_29988732
Cept1:X2_209143856_209152642
B
Homer1:22950023-22964734:+
Slc8a1:3807346-4520635:-
Slc8a1:4259110-4315682:-
Slc8a1:4298605-4315682:-
Slc8a1:4518806-4520635:-
circular/(circ+linear)
cellbody
dendrites
compartment
treatment
control
ptx
C
Homer1:22950023-22964734:+
Slc8a1:3807346-4520635:-
Slc8a1:4259110-4315682:-
Slc8a1:4298605-4315682:-
Slc8a1:4518806-4520635:-
loclogcpm.circ
cellbody
dendrites
compartment
treatment
control
ptx
D
Homer1:22950023-22964734:+
Slc8a1:3807346-4520635:-
Slc8a1:4259110-4315682:-
Slc8a1:4298605-4315682:-
Slc8a1:4518806-4520635:-
loclogcpm.linear
cellbody
dendrites
compartment
treatment
control
ptx

### Suppl. Fig 3: Ptx-dependent impacts on compartmentalization of circRNAs

A. Heatmap of the top 35 circRNAs with greatest differential expression upon Ptx treatment of rat hippocampal neurons regardless of compartment. B. Selected circRNAs (circHomer1, circSlc8a1) isoforms which demonstrate Ptx dependent expression changes (FDR<0.1, circ/circ+lin. Ptx effect irrespective of compartment). C. circRNA BSJ counts for circHomer/circSlc8a1 isoforms. D. linear exon counts are shown for circHomer/circSlc8a1 isoforms.

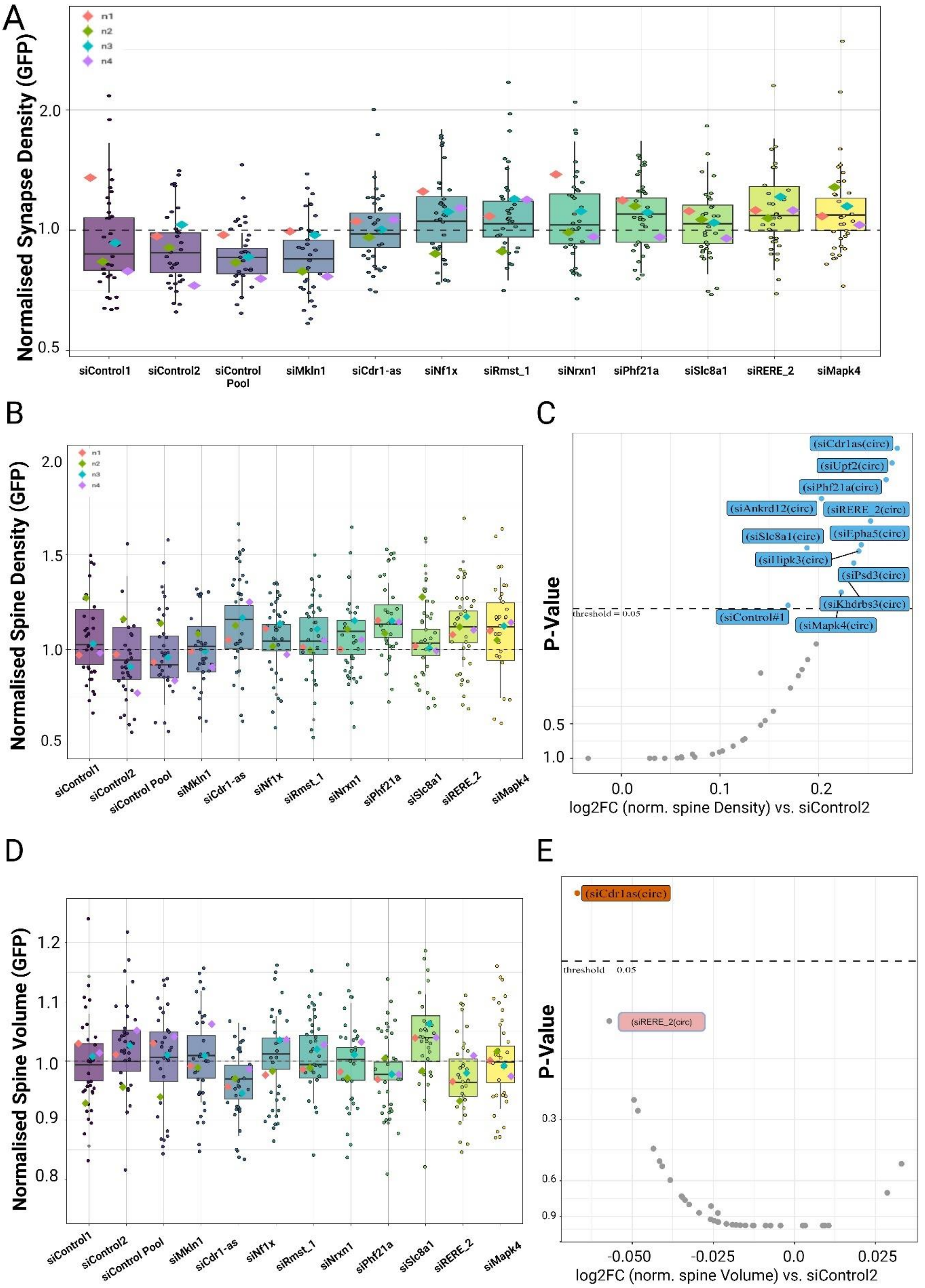
A
n1
n2
n3
n4
Normalised Synapse Density (GFP)
2.0
1.0
0.5
siControl1
siControl2
siControl Pool
siMkln1
siCdr1-as
siNf1x
siRmst_1
siNrxn1
siPhf21a
siSlc8a1
siRERE_2
siMapk4
B
Normalised Spine Density (GFP)
2.0
1.5
1.0
0.5
C
(siCdr1as(circ))
(siUpf2(circ))
(siPhf21a(circ))
(siAnkrd12(circ))
(siRERE_2(circ))
(siEpha5(circ))
(siSlc8a1(circ))
(siPsd3(circ))
(siKhdrbs3(circ))
(siControl#1)
(siMapk4(circ))
threshold = 0.05
P-Value
0.5
1.0
0.0
0.1
0.2
log2FC (norm. spine Density) vs. siControl2
D
Normalised Spine Volume (GFP)
1.2
1.1
1.0
0.9
0.8
E
(siCdr1as(circ))
threshold 0.05
(siRERE_2(circ))
P-Value
0.3
0.6
0.9
-0.050
-0.025
0.0
0.025
log2FC (norm. spine Volume) vs. siControl2

**Suppl. Fig 4: Extended circRNA screen results for synapse co-clusters and dendritic spines.**

A. Box plot related to main Fig. 2C, but showing additional controls (siControl2, siControl_pool). The 7 circRNA siRNAs which demonstrate increased synapse co-cluster density upon knockdown relative to siControl1 also increase relative to siControl2 and siControlPool.

B. Boxplot of circRNA candidate dendritic spine density upon circRNA knockdown with siRNA pools. siControl1 behaves differently to siControl2 and siControl pool.

C. Volcano plot of circRNA candidate dendritic spine density upon circRNA knockdown with siRNA pools. Depicted are 11/32 circRNA candidates with a significantly increased spine density compared to siControl2 normalised to GFP condition, as determined by GLMM statistical modelling. Note siControl1 significantly different to siControl2.

D. Boxplot of circRNA candidate GFP normalized dendritic spine volume upon circRNA knockdown with siRNA pools.

E. Volcano plot of GFP normalized dendritic spine volume relative to siControl2. Depicted is 1 circRNA candidate, Cdr1-as with a significantly decreased spine volume compared to siControl2 normalised to GFP condition, as determined by GLMM statistical modelling.

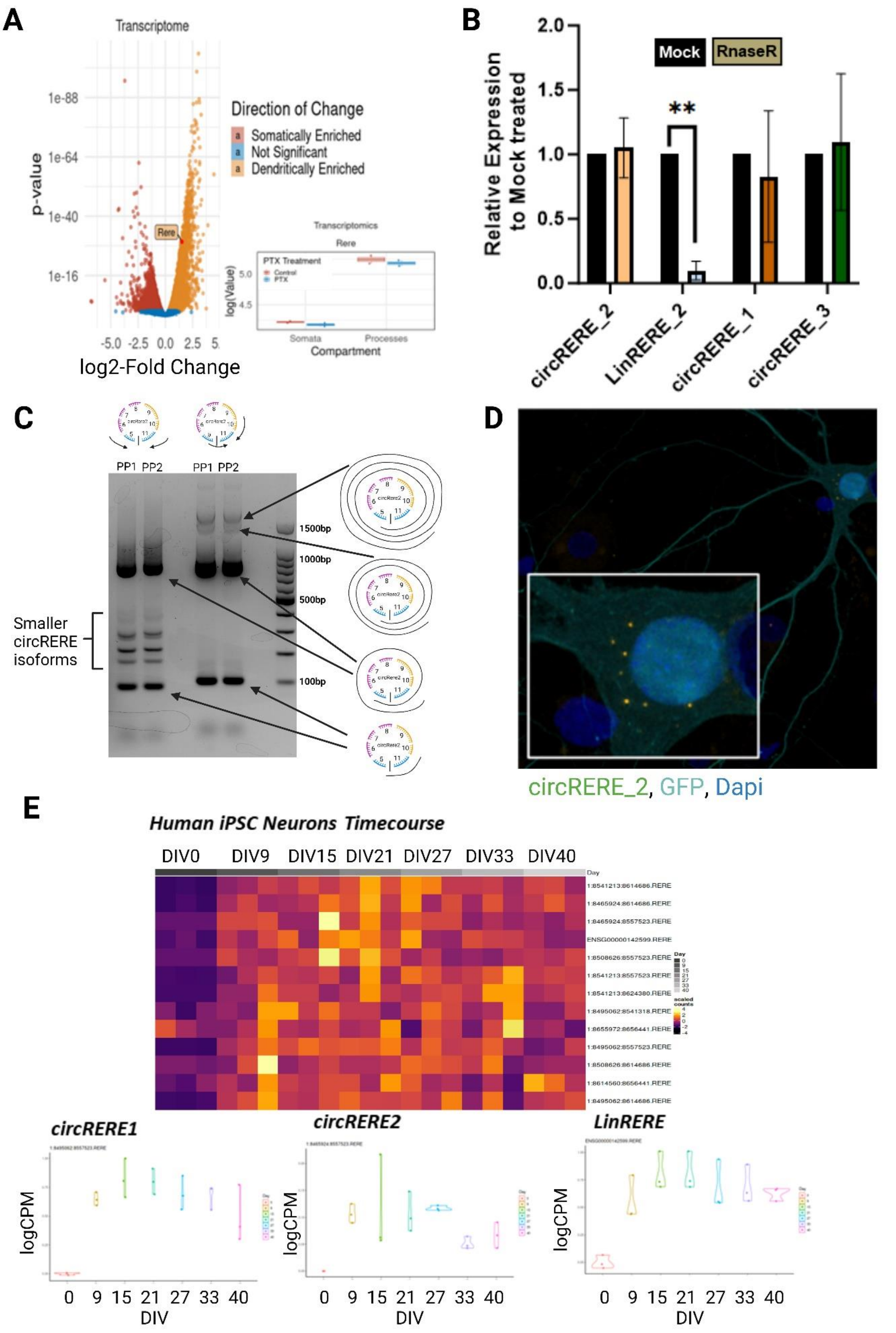
A
Transcriptome
p-value
log2-Fold Change
Direction of Change
Somatically Enriched
Not Significant
Dendritically Enriched
Rere
B
Relative Expression to Mock treated
Mock
RnaseR
**
circRERE_2
LinRERE_2
circRERE_1
circRERE_3
C
PP1 PP2
PP1 PP2
1500bp
1000bp
500bp
100bp
Smaller circRERE isoforms
D
circRERE_2, GFP, Dapi
E
Human iPSC Neurons Timecourse
DIV0 DIV9 DIV15 DIV21 DIV27 DIV33 DIV40
circRERE1
circRERE2
LinRERE
logCPM
DIV

**Supp. Fig 5: Extended circRERE expression analysis.**

A. Volcano plot of all RERE gene reads from Ribo-Minus seq (Colameo et al., 2021), demonstrating overall RERE transcript enrichment in the process compartment. B. qPCR for indicated circular or linear RERE isoforms in either mock or RNAseR-treated extracts from hippocampal neurons. Expression is shown relative to an E.Coli spike-in. (CysG detection). Values obtained from mock treated samples were set to 1.0. N=3 biological replicates. 1-Way ANOVA, $P<0.01$ **. C. Rolling circle amplification of circRERE2. Adult rat cortical RNA was subjected to reverse transcription with random hexamers and PCR-amplified using two different pairs of BSJ-flanking or BSJ-spanning divergent qPCR primers. Running PCR samples on TAE 1.5% Agarose gel electrophoresis revealed the expected size for the respective circRNA species. Flanking qPCR primers also amplify smaller circRERE isoforms due to their nested nature and shared exon identity, whereas qPCR with a BSJ Spanning qPCR primer is specific to circRERE2 alone and demonstrates multiple cycles. D. smFISH in rat hippocampal neurons (DIV15) using a probe directed against the circRERE2 BSJ, illustrating low, exclusive somatic signal likely due to the low affinity of the BSJ-targeting FISH probe. E. Timecourse Ribo-Minus sequencing of all circRERE isoforms and linRERE mRNA throughout the development of human iPSCs (Soutschek et al., 2023).

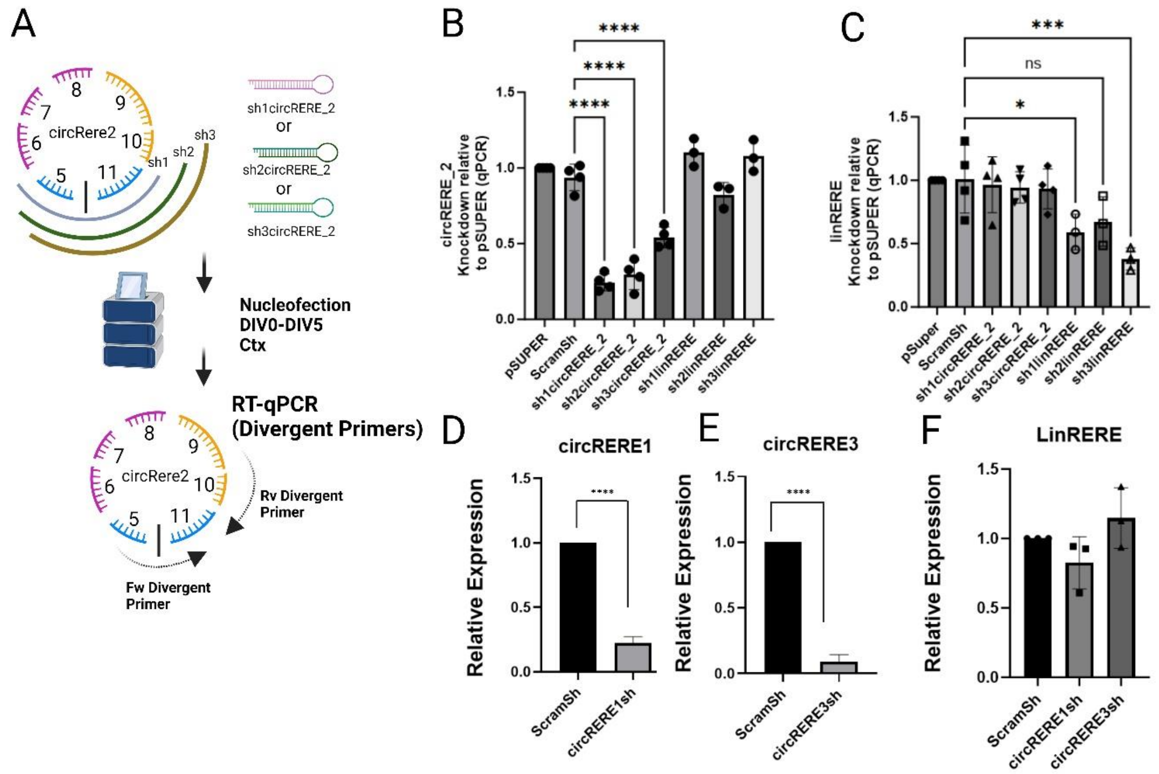

**Supp. Fig 6: Validation of circRERE isoform knockdown efficiency/specificity**

A.Schematic for the shRNA and primer design for circRERE2 RT-qPCR validation. B/D/E. Relative expression levels of indicated circRERE isoforms upon nucleofection/electroporation of the respective shRNA constructs as determined by qPCR using BSJ spanning divergent primers, Ywhaz normalization. C/F. Relative expression levels of linear RERE upon nucleofection of the respective shRNA constructs as determined by qPCR . N=3-4, P<0.05 *, p<0.01** P<0.001 ***, P<0.0001 ****.

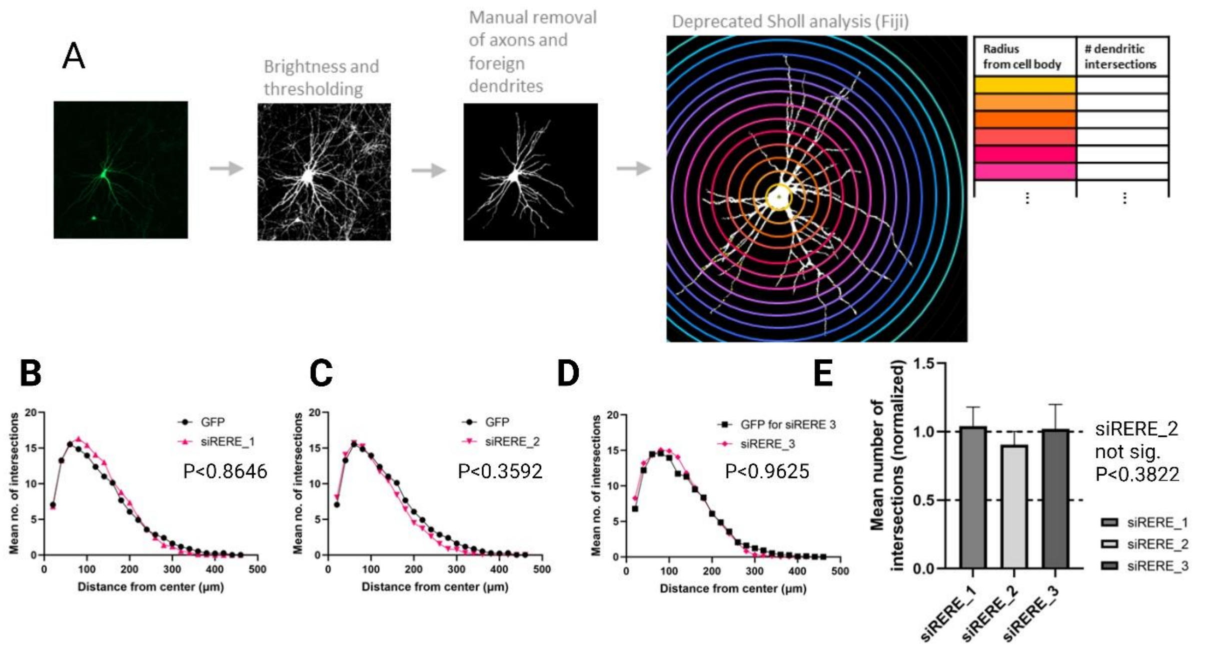


**Supp. Fig 7: Extended morphological analysis of circRERE kd neurons.**

A. Illustration of the Sholl analysis pipeline used for the analysis of dendritogenesis. B-D: Mean number of intersections in rat hippocampal neurons (DIV12) transfected with the indicated siRNAs (red) or control GFP plasmid (black). 1-way ANOVA, Sidak Multiple Comparison test. No significant changes, siRERE_2 (P<0.3592). E. Mean number of total intersections for conditions described on the left. N=4 biological replicates, 8 Cells per conditions, normalised to GFP Control condition. 1-way ANOVA, Sidak Multiple Comparison test. No significant changes Mean intersections (P<0.3822).

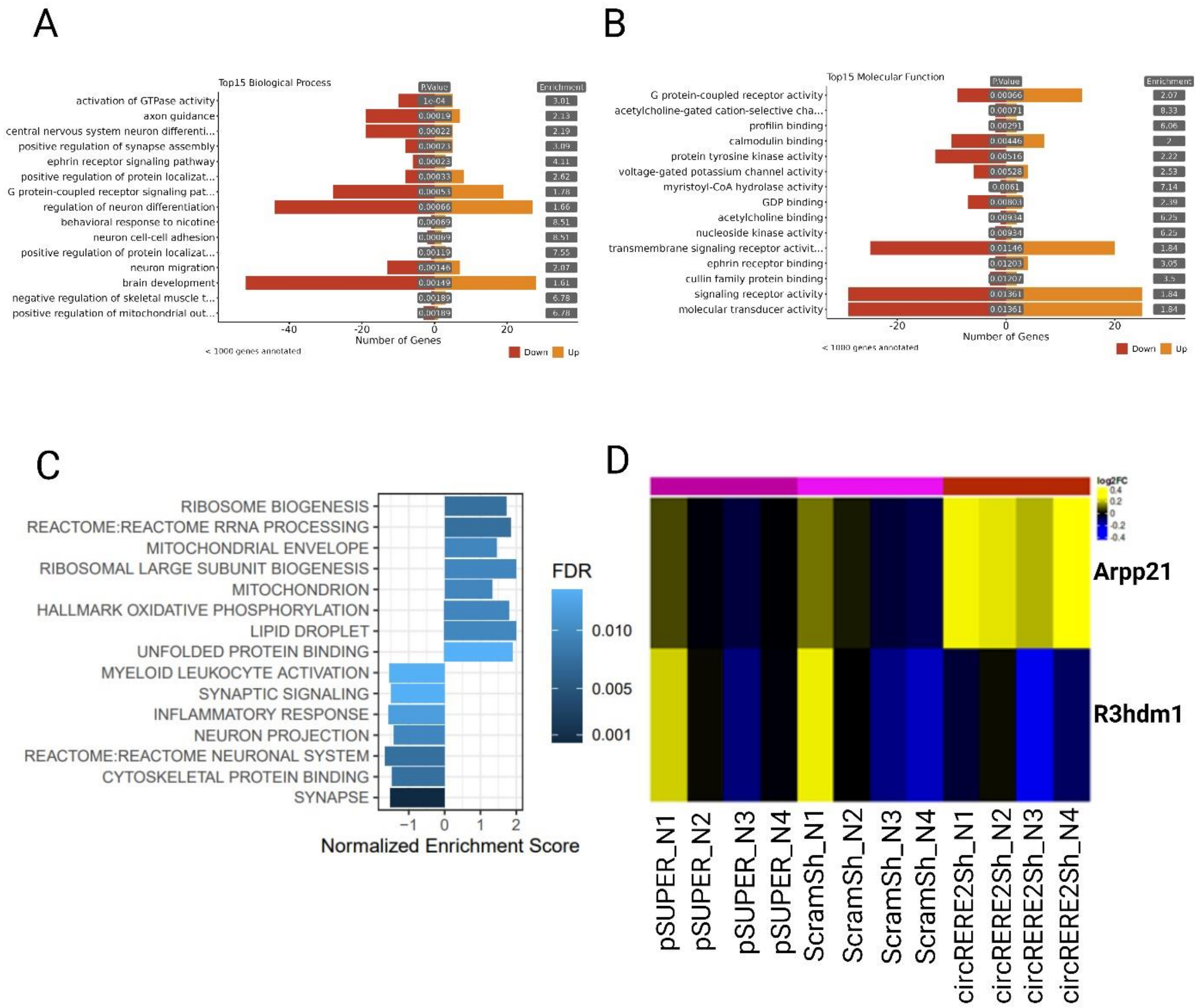


**Supp. Fig 8: circRERE2kd Poly-A seq additional characterisation**

A. Biological Process and B. Molecular Function GO Term analysis (topGO) of DEGs (circRERE2Sh vs.ScramSh/pSUPER Controls). C. Gene-set enrichment analysis in (circRERE2Sh vs.  ScramSh/pSUPER Controls). D. Heatmap of miR-128-3p host genes (Arpp21, R3hdm1) upon circRERE2Sh. Arpp21 expression is (non-significantly) (FDR<0.0674) increased (circRERE2Sh vs.ScramSh/pSUPER Controls).

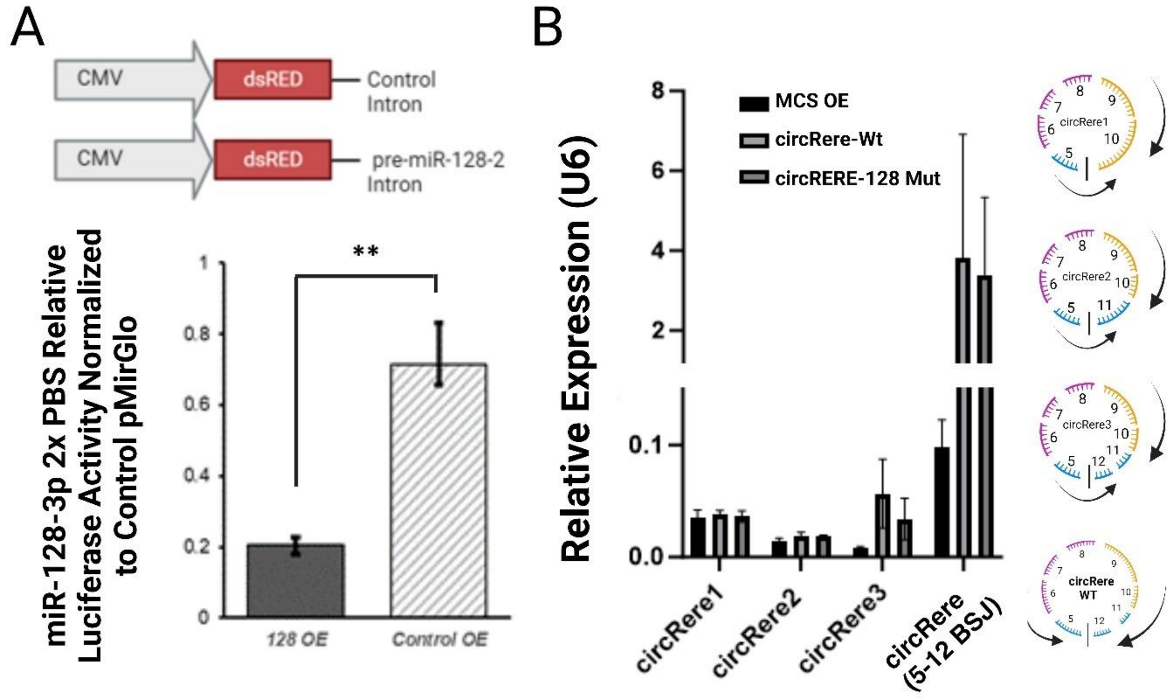


**Supp. Fig 9: Validations of miR-128-3p and circRERE Overexpressions**

A. Upper panel: Illustration of dsRED-pre-miR-128-2 overexpression construct and control. Lower panel: Relative luciferase activity in neurons transfected with miR-128-3p 2x PBS together with the indicated miR-128-3p overexpression (OE) constructs. N=3. Unpaired student’s t-test P<0.01 **. B. Relative expression of indicated circRERE isoforms in primary cortical neurons electroporated with the indicated circRERE or control OE constructs as determined by qPCR. circRERE isoform structure and qPCR primer design is shown on the right. N=3 independent biological replicates, U6 Normalization.

### Supplementary Primer List :

**qPCR primers (Rattus Norvegicus)**

GAPDH fw: GCCTTCTCTTGTGACAAAGTGGA

GAPDH rv: CCGTGGGTAGAGTCATACTGGAA

Ywhaz Fw: CTCCGGACACAGAATATCCAGT

Ywhaz Rv: TGCCATGTCATCGTATCGCT

U6 snRNA Fw: CTCGCTTCGGCAGCACA

U6 snRNA Rv: AACGCTTCACGAATTTGCGT

Cdr1-as (Circular) Fw: CTGCCGTATCCAGGGGTTT

Cdr1-as (Circular) Rv: TGGAAGACCTTGACAGTGTTGG

circRims2 Fw: CTCTCACGGAAAAGTCGCAGT

circRims2 Rv: AGAGGCCGTTGTTCTGTTGATC

linRims2Tr1 Fw: GCGTAGGTGAGAAAGGAGACA

linRims2Tr1 Rv: CTGCCAGGTCACAGGGT

circHomer1 Fw: TGCCATTTTCACATAGGGAAC

circHomer1 Rv: TAACTGCATGCTTGCTGGTG

circGigyf2 Fw: AGGAAGAAAAGATGTAGGCTCCG

circGigyf2 Rv: TCGGCCATATCGATAATCTGC

linGigyf2 Fw: ACAGCCTGCAGTTTGGGAA

linGigyf2 Rv: AGCTTTGGCCTTTTCCAGC

circNf1x Fw: GACCTTTATCTGGCTTACTTTGTCC

circNf1x Rv: GAACCAGGTGTAGGAGAAGGC

linNf1x Fw: CACACCCAACCATCCGCTAC

linNf1x Rv: TTGTGAATGCTGCCCGGT

circStau2 Fw: CAACTTCCGGGGCATGTA

circStau2 Rv: GGGAGACCTGGAGAGAAGCTG

linStau2 Fw: TGAAGTTGCGACTGGAACAG

linStau2 Rv: TGATCCTGAAGACTAGTGGA

circSlc8a1 Fw: CCAGAATGATGAAATAGTGTTGGAAC

circSlc8a1 Rv:  CCACCAGAGTTACCAGACGAAAT

circAnks1b: Fw GGAAGCCAGAGTGTAACAGAGAAG

circAnks1b: Rv CCATAGAGAATCATGATAGCTACCTGT

circSnap25 Fw: GACACCCAGAATCGCCAGA

circSnap25 Rv: CCAGCATCTTTGTTGCACG

circPsd3 Fw: CTCCAAGGATCTTCTAAAATAGTGGC

circPsd3 Rv: CTCTGTGGTCTCCTTTTCCAGAA

circAnks1b Fw: GGAAGCCAGAGTGTAAACAACG

circAnks1b Rv: GGGATCGGTGAGCTCCTCTA

circNckap1 Fw: TTAACGAGCTTGTGCAGAATCTCT

circNckap1 Rv: GCTTTCTTACCTTGTTTCACACTTAGG

CysG Fw E.Coli: TTGTCGGCGGTGGTGATGTC

CysG Rv E.Coli: ATGCGGTGAACTGTGGAATAAACG

**circRERE2 Flanking (2 sets, Rolling Circle Amp)**

circRERE2 Flanking fw1: GTGCCCAAGCTGATCGAGAA

circRERE2 Flanking Rv1: CTGACGGTAGTACCATTTGACG

circRERE2 Flanking Fw2: TCGAGAAGTGCTGGACAGAG

circRERE2 Flanking Rv2: CTGGAACTTCAGACTGACGGTAG

**circRERE2 BSJ spanning (qPCR)**

circRERE2 BSJ spanning Fw: TGGACAGAGGATGAAGTGAGT

circRERE2 BSJ spanning Rv: GGTCCTGAACCAAATGCTGA

**circRERE1 BSJ Spanning (qPCR)**

circRERE1 BSJ Spanning Fw: GCACTGAACACAAGTAAGAGG

circRERE1 BSJ Spanning Rv: TGCCGGTCCTGAACCAAATG

**circRERE3 BSJ Spanning (qPCR)**

circRERE3 BSJ Spanning Fw: AGGAAACCAGTAAGAGGGACCA

circRERE3 BSJ Spanning Rv: ATGCCGGTCCTGAACCAAAT

**linRERE (qPCR)**

linRERE Fw: CCTCACTTAGCTCGCTTCCC

linRERE Rv: GGTAGGGGGTGCCAAAAACT

**siRNAs:**

| **Target RNA Sequence / siRNA ID** | **Primer Name** | **sense** | **antisense** |
|---|---|---|---|
| ACUUAAAGUGUUCUAAGAA | **Whsc1 #1** | **ACUUAAAGUGUUCUAAGAATT** | **UUCUUAGAACACUUUAAGUTT** |
| AAACUUAAAGUGUUCUAAG | **Whsc1 #2** | **AAACUUAAAGUGUUCUAAGTT** | **CUUAGAACACUUUAAGUUUTT** |
| CCAAACUUAAAGUGUUCUA | **Whsc1 #3** | **CCAAACUUAAAGUGUUCUATT** | **UAGAACACUUUAAGUUUGGTT** |
| GGUAAAUGGAACAUUUUAG | **Mkln1 #1** | **GGUAAAUGGAACAUUUUAGTT** | **CUAAAAUGUUCCAUUUACCTT** |
| GCGGUAAAUGGAACAUUUU | **Mkln1 #2** | **GCGGUAAAUGGAACAUUUUTT** | **AAAAUGUUCCAUUUACCGCTT** |
| AAGCGGUAAAUGGAACAUU | **Mkln1 #3** | **AAGCGGUAAAUGGAACAUUTT** | **AAUGUUCCAUUUACCGCUUTT** |
| GAAAUAGUGUUGGAACAAU | **Slc8a1 #1** | **GAAAUAGUGUUGGAACAAUTT** | **AUUGUUCCAACACUAUUUCTT** |
| AUGAAAUAGUGUUGGAACA | **Slc8a1 #2** | **AUGAAAUAGUGUUGGAACATT** | **UGUUCCAACACUAUUUCAUTT** |
| UGAUGAAAUAGUGUUGGAA | **Slc8a1 #3** | **UGAUGAAAUAGUGUUGGAATT** | **UUCCAACACUAUUUCAUCATT** |
| UUAUACAGAUCCAGGAUGA | **Ankrd12 #1** | **UUAUACAGAUCCAGGAUGATT** | **UCAUCCUGGAUCUGUAUAATT** |
| AGUUAUACAGAUCCAGGAU | **Ankrd12 #2** | **AGUUAUACAGAUCCAGGAUTT** | **AUCCUGGAUCUGUAUAACUTT** |
| AAAGUUAUACAGAUCCAGG | **Ankrd12 #3** | **AAAGUUAUACAGAUCCAGGTT** | **CCUGGAUCUGUAUAACUUUTT** |
| CAAUACAGUGCAAUGGCAA | **Nrcam #1** | **CAAUACAGUGCAAUGGCAATT** | **UUGCCAUUGCACUGUAUUGTT** |
| ACAAUACAGUGCAAUGGCA | **Nrcam #2** | **ACAAUACAGUGCAAUGGCATT** | **UGCCAUUGCACUGUAUUGUTT** |
| AUACAGUGCAAUGGCAAGU | **Nrcam #3** | **AUACAGUGCAAUGGCAAGUTT** | **ACUUGCCAUUGCACUGUAUTT** |
| CAAUGACCGCAAAGGUCCU | **Nrxn1 #1** | **CAAUGACCGCAAAGGUCCUTT** | **AGGACCUUUGCGGUCAUUGTT** |
| UGCAAUGACCGCAAAGGUC | **Nrxn1 #2** | **UGCAAUGACCGCAAAGGUCTT** | **GACCUUUGCGGUCAUUGCATT** |
| UCUGCAAUGACCGCAAAGG | **Nrxn1 #3** | **UCUGCAAUGACCGCAAAGGTT** | **CCUUUGCGGUCAUUGCAGATT** |
| UGUCAAGUGGGAAGAGAUU | **Epha5 #1** | **UGUCAAGUGGGAAGAGAUUTT** | **AAUCUCUUCCCACUUGACATT** |

| | | | |
|---|---|---|---|
| CUGUCAAGUGGGA AGAGAU | **Epha5 #2** | **CUGUCAAGUGGGA AGAGAUTT** | **AUCUCUUCCCACU UGACAGTT** |
| CCUGUCAAGUGGG AAGAGA | **Epha5 #3** | **CCUGUCAAGUGGG AAGAGATT** | **UCUCUUCCCACUU GACAGGTT** |
| CCAAAGAUACUCA UGCUAA | **Rmst_1 #1** | **CCAAAGAUACUCA UGCUAATT** | **UUAGCAUGAGUAU CUUUGGTT** |
| GCCAAAGAUACUC AUGCUA | **Rmst_1 #2** | **GCCAAAGAUACUC AUGCUATT** | **UAGCAUGAGUAUC UUUGGCTT** |
| AGCCAAAGAUACU CAUGCU | **Rmst_1 #3** | **AGCCAAAGAUACU CAUGCUTT** | **AGCAUGAGUAUCU UUGGCUTT** |
| CACACUCCGGGAU GAGUUC | **Nf1x #1** | **CACACUCCGGGAU GAGUUCTT** | **GAACUCAUCCCGG AGUGUGTT** |
| CUCCGGGAUGAGU UCCACC | **Nf1x #2** | **CUCCGGGAUGAGU UCCACCTT** | **GGUGGAACUCAUC CCGGAGTT** |
| UCCACACUCCGGG AUGAGU | **Nf1x #3** | **UCCACACUCCGGG AUGAGUTT** | **ACUCAUCCCGGAG UGUGGATT** |
| UGAACACAAGUAA GAGGGA | **RERE_1 #1** | **UGAACACAAGUAA GAGGGATT** | **UCCCUCUUACUUG UGUUCATT** |
| GAACACAAGUAAG AGGGAC | **RERE_1 #2** | **GAACACAAGUAAG AGGGACTT** | **GUCCCUCUUACUU GUGUUCTT** |
| AACACAAGUAAGA GGGACC | **RERE_1 #3** | **AACACAAGUAAGA GGGACCTT** | **GGUCCCUCUUACU UGUGUUTT** |
| CCAGAGUGUAACA GAGAAG | **Anks1b_1 #1** | **CCAGAGUGUAACA GAGAAGTT** | **CUUCUCUGUUACA CUCUGGTT** |
| GCCAGAGUGUAAC AGAGAA | **Anks1b_1 #2** | **GCCAGAGUGUAAC AGAGAATT** | **UUCUCUGUUACAC UCUGGCTT** |
| AGUGUAACAGAGA AGGGGA | **Anks1b_1 #3** | **AGUGUAACAGAGA AGGGGATT** | **UCCCCUUCUCUGU UACACUTT** |
| AGAGUGUAAACAA CGAGAA | **Anks1b_2 #1** | **AGAGUGUAAACAA CGAGAATT** | **UUCUCGUUGUUUA CACUCUTT** |
| GCCAGAGUGUAAA CAACGA | **Anks1b_2 #2** | **GCCAGAGUGUAAA CAACGATT** | **UCGUUGUUUACAC UCUGGCTT** |
| GAGUGUAAACAAC GAGAAC | **Anks1b_2 #3** | **GAGUGUAAACAAC GAGAACTT** | **GUUCUCGUUGUUU ACACUCTT** |
| AUGAAGUGAGUAA GAGGGA | **RERE_2 #1** | **AUGAAGUGAGUAA GAGGGATT** | **UCCCUCUUACUCA CUUCAUTT** |
| GAAGUGAGUAAGA GGGACC | **RERE_2 #2** | **GAAGUGAGUAAGA GGGACCTT** | **GGUCCCUCUUACU CACUUCTT** |
| GGAUGAAGUGAGU AAGAGG | **RERE_2 #3** | **GGAUGAAGUGAGU AAGAGGTT** | **CCUCUUACUCACU UCAUCCTT** |
| GGCAAGAAAUAUA GGAGUA | **Foxn2 #1** | **GGCAAGAAAUAUA GGAGUATT** | **UACUCCUAUAUUU CUUGCCTT** |
| CGGCAAGAAAUAU AGGAGU | **Foxn2 #2** | **CGGCAAGAAAUAU AGGAGUTT** | **ACUCCUAUAUUUC UUGCCGTT** |
| ACGGCAAGAAAUA UAGGAG | **Foxn2 #3** | **ACGGCAAGAAAUA UAGGAGTT** | **CUCCUAUAUUUCU UGCCGUTT** |
| UUUCACAUAGGGA ACAACC | **Homer1 #1** | **UUUCACAUAGGGA ACAACCTT** | **GGUUGUUCCCUAU GUGAAATT** |
| CCAUUUUCACAUA GGGAAC | **Homer1 #2** | **CCAUUUUCACAUA GGGAACTT** | **GUUCCCUAUGUGA AAAUGGTT** |

| | | | |
|---|---|---|---|
| UCACAUAGGGAAC AACCUA | **Homer1 #3** | **UCACAUAGGGAAC AACCUATT** | **UAGGUUGUUCCCU AUGUGATT** |
| UUGCUGCGAAUUU UAUAGG | **Uvrag #1** | **UUGCUGCGAAUUU UAUAGGTT** | **CCUAUAAAAUUCG CAGCAATT** |
| CUUUGCUGCGAAU UUUAUA | **Uvrag #2** | **CUUUGCUGCGAAU UUUAUATT** | **UAUAAAAUUCGCA GCAAAGTT** |
| UUCUUUGCUGCGA AUUUUA | **Uvrag #3** | **UUCUUUGCUGCGA AUUUUATT** | **UAAAAUUCGCAGC AAAGAATT** |
| AUGGAGAGUAUUU CAACUU | **Khdrbs3 #1** | **AUGGAGAGUAUUU CAACUUTT** | **AAGUUGAAAUACU CUCCAUTT** |
| GGAGAGUAUUUCA ACUUUG | **Khdrbs3 #2** | **GGAGAGUAUUUCA ACUUUGTT** | **CAAAGUUGAAAUA CUCUCCTT** |
| GAGAGUAUUUCAA CUUUGU | **Khdrbs3 #3** | **GAGAGUAUUUCAA CUUUGUTT** | **ACAAAGUUGAAAU ACUCUCTT** |
| GCCUGUGUGGGA AUAGCUA | **Zranb1 #1** | **GCCUGUGUGGGAA UAGCUATT** | **UAGCUAUUCCCAC ACAGGCTT** |
| CCUGUGUGGGAAU AGCUAU | **Zranb1 #2** | **CCUGUGUGGGAAU AGCUAUTT** | **AUAGCUAUUCCCA CACAGGTT** |
| AUGCCUGUGUGG GAAUAGC | **Zranb1 #3** | **AUGCCUGUGUGGG AAUAGCTT** | **GCUAUUCCCACAC AGGCAUTT** |
| UGUGCAAAAAGAA ACUUGU | **Ubn2 #1** | **UGUGCAAAAAGAA ACUUGUTT** | **ACAAGUUUCUUUU UGCACATT** |
| UCUGUGCAAAAAG AAACUU | **Ubn2 #2** | **UCUGUGCAAAAAG AAACUUTT** | **AAGUUUCUUUUUG CACAGATT** |
| CUUCUGUGCAAAA AGAAAC | **Ubn2 #3** | **CUUCUGUGCAAAA AGAAACTT** | **GUUUCUUUUUGCA CAGAAGTT** |
| AUGGCUUCAUGAG AUACUC | **Rmst_2 #1** | **AUGGCUUCAUGAG AUACUCTT** | **GAGUAUCUCAUGA AGCCAUTT** |
| GGCUUCAUGAGAU ACUCAU | **Rmst_2 #2** | **GGCUUCAUGAGAU ACUCAUTT** | **AUGAGUAUCUCAU GAAGCCTT** |
| UCAUGAGAUACUC AUGCUA | **Rmst_2 #3** | **UCAUGAGAUACUC AUGCUATT** | **UAGCAUGAGUAUC UCAUGATT** |
| AGGAAACCAGUAA GAGGGA | **RERE_3 #1** | **AGGAAACCAGUAA GAGGGATT** | **UCCCUCUUACUGG UUUCCUTT** |
| GAAACCAGUAAGA GGGACC | **RERE_3 #2** | **GAAACCAGUAAGA GGGACCTT** | **GGUCCCUCUUACU GGUUUCTT** |
| UAAGGAAACCAGU AAGAGG | **RERE_3 #3** | **UAAGGAAACCAGU AAGAGGTT** | **CCUCUUACUGGUU UCCUUATT** |
| AUGGCAAGGUAGG UGAGCC | **Mapkap1 #1** | **AUGGCAAGGUAGG UGAGCCTT** | **GGCUCACCUACCU UGCCAUTT** |
| UGAUGGCAAGGUA GGUGAG | **Mapkap1 #2** | **UGAUGGCAAGGUA GGUGAGTT** | **CUCACCUACCUUG CCAUCATT** |
| UUUGAUGGCAAGG UAGGUG | **Mapkap1 #3** | **UUUGAUGGCAAGG UAGGUGTT** | **CACCUACCUUGCC AUCAAATT** |
| UCAUGUUGGUUCC CAGUCA | **Ash1l #1** | **UCAUGUUGGUUCC CAGUCATT** | **UGACUGGGAACCA ACAUGATT** |
| CAUUCAUGUUGGU UCCCAG | **Ash1l #2** | **CAUUCAUGUUGGU UCCCAGTT** | **CUGGGAACCAACA UGAAUGTT** |
| AUUCAUGUUGGUU CCCAGU | **Ash1l #3** | **AUUCAUGUUGGUU CCCAGUTT** | **ACUGGGAACCAAC AUGAAUTT** |

| | | | |
|---|---|---|---|
| AGAAACAGGGGAC UAGAAA | **Phf21a #1** | **AGAAACAGGGGAC UAGAAATT** | **UUUCUAGUCCCCU GUUUCUTT** |
| GAGAAACAGGGGA CUAGAA | **Phf21a #2** | **GAGAAACAGGGGA CUAGAATT** | **UUCUAGUCCCCUG UUUCUCTT** |
| AAACAGGGGACUA GAAAGC | **Phf21a #3** | **AAACAGGGGACUA GAAAGCTT** | **GCUUUCUAGUCCC CUGUUUTT** |
| AGGCAGUGUGAAA ACUCAG | **Ralgapa1 #1** | **AGGCAGUGUGAAA ACUCAGTT** | **CUGAGUUUUCACA CUGCCUTT** |
| CAAGGCAGUGUGA AAACUC | **Ralgapa1 #2** | **CAAGGCAGUGUGA AAACUCTT** | **GAGUUUUCACACU GCCUUGTT** |
| UUCAAGGCAGUGU GAAAAC | **Ralgapa1 #3** | **UUCAAGGCAGUGU GAAAACTT** | **GUUUUCACACUGC CUUGAATT** |
| CCACAAGGGAAGG GGACUG | **Mapk4 #1** | **CCACAAGGGAAGG GGACUGTT** | **CAGUCCCCUUCCC UUGUGGTT** |
| CCCACAAGGGAAG GGGACU | **Mapk4 #2** | **CCCACAAGGGAAG GGGACUTT** | **AGUCCCCUUCCCU UGUGGGTT** |
| CUCCCACAAGGGA AGGGGA | **Mapk4 #3** | **CUCCCACAAGGGA AGGGGATT** | **UCCCCUUCCCUUG UGGGAGTT** |
| UUCUAAAAUAGUG GCUAUU | **Psd3 #1** | **UUCUAAAAUAGUG GCUAUUTT** | **AAUAGCCACUAUU UUAGAATT** |
| UCUUCUAAAAUAG UGGCUA | **Psd3 #2** | **UCUUCUAAAAUAG UGGCUATT** | **UAGCCACUAUUUU AGAAGATT** |
| GAUCUUCUAAAAU AGUGGC | **Psd3 #3** | **GAUCUUCUAAAAU AGUGGCTT** | **GCCACUAUUUUAG AAGAUCTT** |
| AGAUGUAGGCUCC GUGCUC | **Gigyf2 #1** | **AGAUGUAGGCUCC GUGCUCTT** | **GAGCACGGAGCCU ACAUCUTT** |
| GAUGUAGGCUCCG UGCUCU | **Gigyf2 #2** | **GAUGUAGGCUCCG UGCUCUTT** | **AGAGCACGGAGCC UACAUCTT** |
| AUGUAGGCUCCGU GCUCUG | **Gigyf2 #3** | **AUGUAGGCUCCGU GCUCUGTT** | **CAGAGCACGGAGC CUACAUTT** |
| CUCAGCAGGUUUG GGCCCC | **pwwp2a #1** | **CUCAGCAGGUUUG GGCCCCTT** | **GGGGCCCAAACCU GCUGAGTT** |
| CGCUCAGCAGGUU UGGGCC | **pwwp2a #2** | **CGCUCAGCAGGUU UGGGCCTT** | **GGCCCAAACCUGC UGAGCGTT** |
| CCCGCUCAGCAGG UUUGGG | **pwwp2a #3** | **CCCGCUCAGCAGG UUUGGGTT** | **CCCAAACCUGCUG AGCGGGTT** |
| GCAAAGGCGUAUU CUGCAC | **Upf2 #1** | **GCAAAGGCGUAUU CUGCACTT** | **GUGCAGAAUACGC CUUUGCTT** |
| AGGCAAAGGCGUA UUCUGC | **Upf2 #2** | **AGGCAAAGGCGUA UUCUGCTT** | **GCAGAAUACGCCU UUGCCUTT** |
| GGCAAAGGCGUAU UCUGCA | **Upf2 #3** | **GGCAAAGGCGUAU UCUGCATT** | **UGCAGAAUACGCC UUUGCCTT** |
| UGCCGUAUCCAGG GGUUUC | **Cdr1as #1** | **UGCCGUAUCCAGG GGUUUCTT** | **GAAACCCCUGGAU ACGGCATT** |
| CCGUAUCCAGGGG UUUCCA | **Cdr1as #2** | **CCGUAUCCAGGGG UUUCCATT** | **UGGAAACCCCUGG AUACGGTT** |
| GUAUCCAGGGGUU UCCAGU | **Cdr1as #3** | **GUAUCCAGGGGUU UCCAGUTT** | **ACUGGAAACCCCU GGAUACTT** |
| CUAUAGGUAUGGC CUCACA | **Hipk3 #1** | **CUAUAGGUAUGGC CUCACATT** | **UGUGAGGCCAUAC CUAUAGTT** |

| | | | |
|---|---|---|---|
| GGUACUAUAGGUA UGGCCU | **Hipk3 #2** | **GGUACUAUAGGUA UGGCCUTT** | **AGGCCAUACCUAU AGUACCTT** |
| UACUAUAGGUAUG GCCUCA | **Hipk3 #3** | **UACUAUAGGUAUG GCCUCATT** | **UGAGGCCAUACCU AUAGUATT** |

Silencer Select Negative Control #1 (Catalog number 4390843, Thermo Fisher), Silencer Select Negative Control#2 (Catalog number 4390847, Thermo Fisher) and a mixture of 50/50 “siControl Pool”, were utilised in the siRNA knockdown screen. These are referred to as siControl1, siControl2 and siControl Pool in the text.

**shRNAs:**

**“pSUPER”** plasmid utilised as a control, all nucleotides excised between BglII and HindII RE sites, preventing shRNA production.

For additional shRNAs below are listed sense sequences:

**ScramSh**: AAACCTTGTGGTCCTTAGG

**linRERE**

sh1 RERE: GGAACGAGAGCGAGAGAAA (RERE exon19)

sh2 RERE: CCAAGAAAGTGAAGGAAGA (RERE exon16-17)

sh3 RERE: ACAGAAAGCCCGAGAGGAA (RERE exon 19)

**circRERE2**

Sh1: ATGAAGTGAGTAAGAGGGA

Sh2: GAAGTGAGTAAGAGGGACC

Sh3: GGATGAAGTGAGTAAGAGG

Note: circRERE2Sh2 noted above was utilised for all experiments after validation of efficacy and specificity.

**circRERE1 Sh**

AACACAAGUAAGAGGGACC

**circRERE3 Sh**

AGGAAACCAGUAAGAGGGA

**circRNA FISH Probes**

**Target sequences for probe design (rat)**
circHomer1: 5’-UUUCACAUAGGGAACAACCU-3’
circRmst_2: 5’-GGCUUCAUGAGAUACUCAUG-3’

circStau2_1: 5’- ACAACCAGAGUUUUUUGGAG -3’

circRERE_2: 5’- GGAUGAAGUGAGUAAGAGGG -3’

**miR-128-3p Perfect binding site reporter insert**

gctagcAAAGAGACCGGTTCACTGTGActAAAGAGACCGGTTCACTGTGgtcgac

Control Reporter possessed no insert between NheI/SalI RE sites.
